# Time in allopatry does not predict the outcome of secondary contact in lowland Panamanian birds

**DOI:** 10.1101/2022.10.25.513737

**Authors:** Jessica F. Mclaughlin, Matthew J. Miller

**Author notes:** Corresponding author: J. F. McLaughlin, 217 Hilgard Hall, University of California Berkeley, Berkeley, CA, USA, 94720. Statement of authorship*: JFM conceived the study in consultation with MJM. JFM generated genomic libraries to supplement the original dataset, conducted all analyses, created original illustrations, and wrote the initial draft, with feedback from MJM on subsequent drafts.

## Abstract

Geographic speciation models assume that time in allopatry should result in greater reproductive isolation between populations. Here we test the prediction that greater time in allopatry results in greater reproductive isolation using comparative ultraconserved element (UCE) data from ten bird lineages in secondary contact in Panama, measuring both genome-wide divergence and the geographic extent of hybridization. The best-fit models for the proportion of fixed Z-linked and autosomal loci to our data includes a combination of both time (as measured by mtDNA divergence) and hand-wing index, emphasizing that the role of time is tempered by dispersal capability. Furthermore, time does not predict the extent of genome-wide introgression as measured by the median width of diagnostic loci clines or the degree of variation in cline centers or widths. These metrics of the outcome of secondary contact were best predicted by ecological and genomic factors, including diet, hand-wing index, and genome-wide *F_ST_* respectively, that are understood to serve as proxies for dispersal, the variability of population size, and overall genomic divergence. We find a primary role for ecological factors instead of isolation time in determining secondary contact outcomes for a lineage, highlighting how ecology shapes the development of reproductive isolation.

## Introduction

Historically, animal speciation has been understood to be driven in its early stages by population isolation in allopatry (1–8). Mayr (1963) outlined a basic model of allopatric speciation in which an ancestral species is divided into two allopatric populations by vicariance or dispersal, leading to isolation without gene flow (2). During time in isolation, differences between the populations – both phenotypic and genetic – accumulate. Under Mayr’s model, one or both populations experience range expansion, and the populations come into secondary contact, where speciation is completed (although later work would regard this as not necessarily required). Later authors would expand on this, describing the outcome of secondary contact as a continuum (9–13), from population fusion – also called reticulation or reverse speciation (14–17) – to complete reproductive isolation (RI) without introgression (18, 19). Intermediate along this continuum is the formation of a hybrid zone, with partial RI (20). This current paradigm for animal speciation is the result of research demonstrating the pervasiveness of gene flow during the speciation process (21–27).

Implicit in both Mayr’s model and more modern interpretations that allow for gene flow is a central role for time in allopatry for driving differentiation and, eventually, speciation. As time increases, the likelihood of evolutionary changes occurring increases in parallel. Time allows for directional selection due to local environmental conditions as well as stochastic evolution due to drift (28, 29). Orr and Turelli (2001) described the ‘speciation clock’ as the outcome of molecular evolution coupled with the accumulation of Dobzhansky-Muller incompatibilities (30). Genomic variations that arise and become fixed, such as chromosomal inversions (31), the mutations within them (32), or mitonuclear incompatibilities (33, 34), likely occur at a largely consistent rate for similarly sized populations (35), but as time in isolation passes, it becomes more likely that any one or more of them occur and accumulate. Over time, this process increases the likelihood of such changes leading to the development of RI. Experimental evidence demonstrates hybrid viability decreasing over time (29, 36–40).

Therefore, time should predict where a lineage is in the process of speciation, and the expected extent of introgression in secondary contact. There is some evidence from birds that supports this hypothesis. Studies of avian hybridization have shown a relationship between time (measured by mitochondrial divergence) and the development of RI (41, 42), with hybrid infertility and inviability becoming more likely with increased divergence time (37, 38). Likewise, traits linked with prezygotic isolation also show this tendency (43, 44), which is relevant given the general pattern in birds of prezygotic isolation arising before postzygotic (41, 45). From these observations, it is reasonable to predict that as time in isolation increases, so will RI from both pre- and postzygotic mechanisms. Thus, in the context of secondary contact, the extent of introgression will decrease in tandem.

However, many factors could decouple time from the outcome of secondary contact and the development of RI. Birds, which tend to have higher rates of hybridization than many vertebrates (41, 46), have been a useful group to study these factors. However, in many cases it has proved difficult to detect a reduction in fitness of hybrids in wild populations, and introgression even appears to be positively selected for in some cases (47; 48). A propensity for hybridization arises from the nature of bird genomes themselves (49). Bird genomes are small compared to other tetrapods (50, 51), likely driven by the metabolic demands of flight (52–54). This is then compounded by the relatively conserved synteny of avian chromosomes (55, 56). As a result of these attributes, there are fewer structural variants which could lead to postzygotic barriers seen in other taxa (56). This likely provides an explanation for the tendency of RI in birds to be maintained by prezygotic barriers (45), but it simultaneously informs why these barriers may be less effective in perpetuating that isolation. Postzygotic barriers are far more effective in limiting hybridization than assortative mating (57), whereas prezygotic barriers can be eroded by environmental changes (58) or even by sexual selection, as seen in birds such as manakins (59–62), fairywrens (63, 64), and jacanas (48), where the phenotype of the courtship-dominant sex in one taxon is actually preferred by the other. These factors all mean that time may not be an effective predictor of the outcome of secondary contact in birds.

We investigated the relationship between time in isolation and the degree of RI using loci linked to ultraconserved elements (UCEs) in ten lineages of birds in secondary contact spanning a range of divergence dates across Panama. We tested several predictions that arise from the time-in-allopatry hypothesis. Firstly, we predict that greater depths of mitochondrial breaks, reflective of longer time in isolation, are correlated with the increased differentiation across the nuclear genome (Figure 1A). We expected that if this is the case, the proportion of loci fixed between the eastern and western Panama populations would be higher in taxa with a greater mitochondrial divergence. Lastly, in older taxa, the geographic extent of admixture will be reduced (Figure 1B), and that geographic clines should be narrower and more consistently located along the transect (Figure 1C).

**Figure 1:**
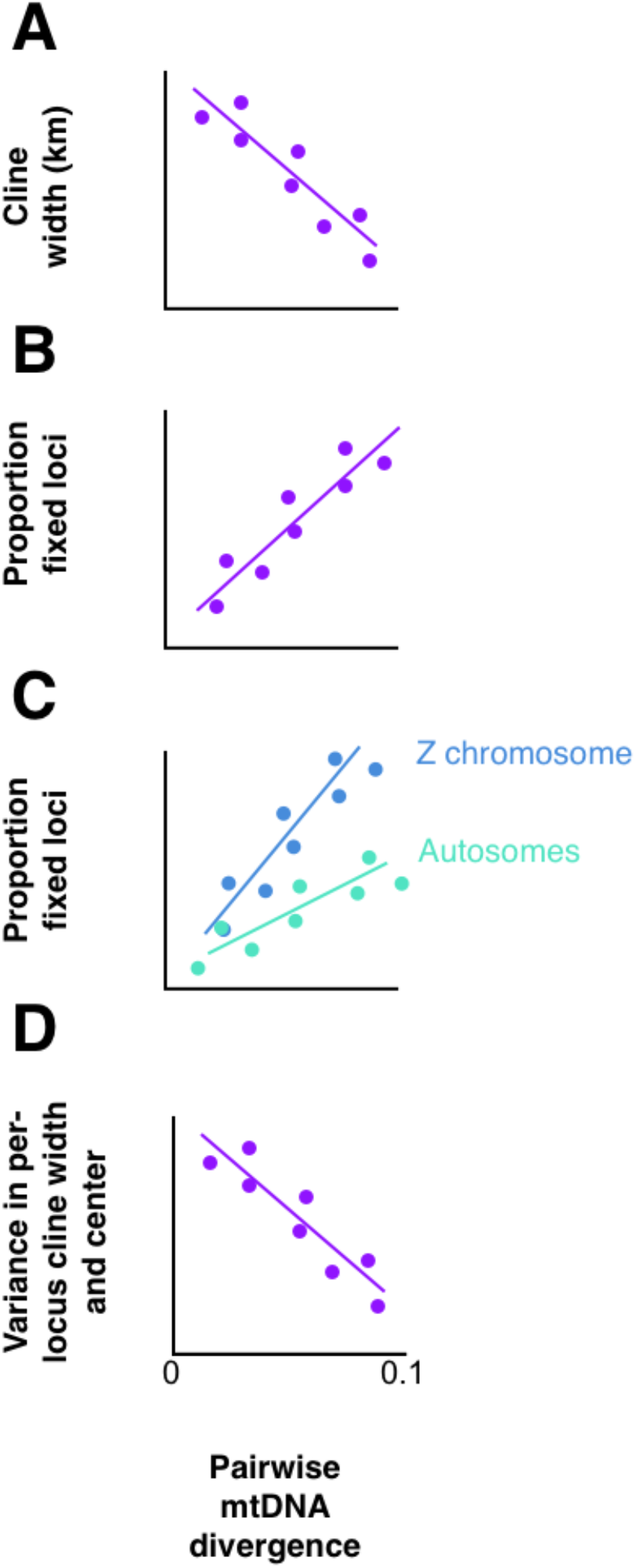
Predictions given time as the main driving factor in determining secondary contact outcomes. According to this hypothesis, A) median cline width should decrease with increasing mitochondrial distance, as hybrid zones narrow; B) the proportion of fixed loci will increase as isolated populations accumulate fixed differences independently of each over, either by drift or by adaptive pressures; C) the Z chromosome, with ¾ the effective population size, will accumulate these differences at approximately twice the rate in autosomes, and D) variance in the location of the cline center and the width across loci will decrease, as the hybrid zone stabilizes and narrows under selection pressures against hybrid offspring.

## Results

### Sequencing results

We sequenced between 0.22 and 6.5 million reads for each UCE-enriched sample and between 40.4 and 229.6 million reads for WGS samples (Table S1). From this, we recovered between 1855 and 4292 loci per taxon, with an average coverage of 39.8× (Table S1). Overall, coverage from non-enriched and enriched libraries was similar, but slightly higher in non-enriched samples (48.2x vs 37.1x). However, several unenriched samples were dropped due to low numbers of recovered UCE loci (between 500-1000 loci), demonstrating that while UCEs can be recovered with ease from WGS reads when circumstances call for such an approach, it is less reliable.

### Mitochondrial divergence and fixation rates

Pairwise divergence of all mitochondrial protein-coding regions ranged from 2.75% (*X. minutus*) to 9.83% (*Schiffornis*) (Table 1). *F_ST_* of UCE-linked loci ranged from 0.265 (*Cy. cyanoides*) and 0.688 (*Schiffornis*), and nucleotide diversity (*π*) between 0.00195 (*P. decurtata*) and 0.00298 (*H. leucosticta*; Table 1). We examined nuclear fixation rates separately between the autosomes and Z-chromosomes for three reasons. Firstly, the effective population size of Z chromosome loci is only ¾ that of autosomal loci. Therefore, under neutral conditions Z chromosome loci will accumulate fixed differences at twice the rate of autosomes (65). Secondly, the Z chromosome has been indicated to disproportionately be the site of loci responsible for the development of reproductive barriers in many studies of avian speciation (49, 66–69), making separate consideration of them important. Finally, Z chromosomes are likely to be the sites of incompatibilities as predicted by Haldane’s rule -- if hybrid fitness is lower in one sex than the other, it will be in the heterogametic sex (8, 41, 70, 71), possibly as incompatibilities arise between the different sex chromosomes (72), between the Z chromosome and autosomes (73), or between the Z chromosome and mitochondrial genomes (74). The total proportion of fixed SNPs in these loci was between 0.0126 (*Ca.nigricapillus*) and 0.226 (*Schiffornis*), with autosomal fixation ranging from 0.0105 (*Ca. nigricapillus*) to 0.220 (*Schiffornis*) and Z-linked fixation ranging from 0.0221 (*P. decurtata*) to 0.480 (*Malacoptila panamensis*; Table 1). While average divergence in insectivores was lower than in birds reliant on plant foods, 4.07% vs 5.92%, the difference was not significant (*t* = −1.50, df = 6.57, *p* = 0.180).

**Table 1:**
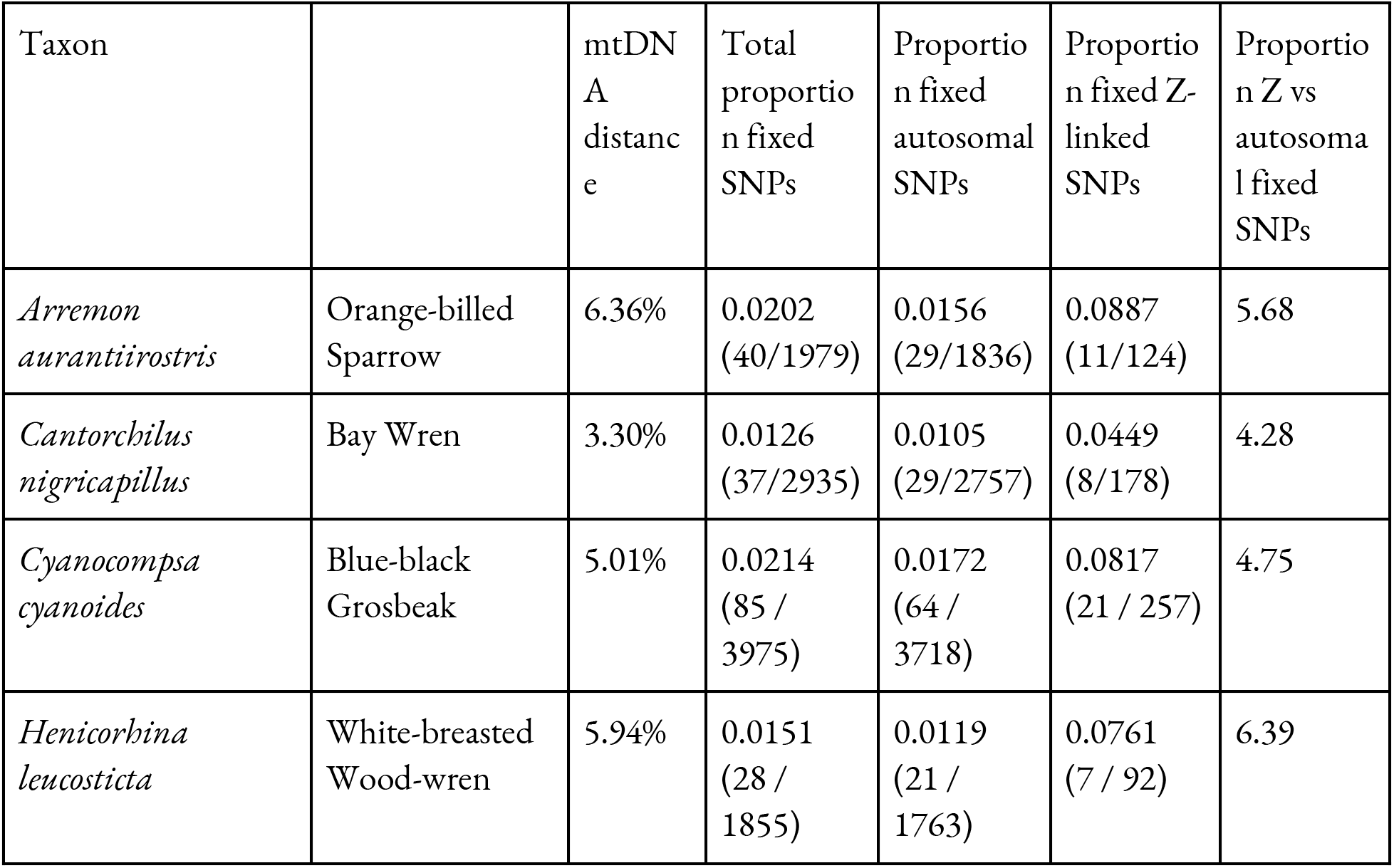
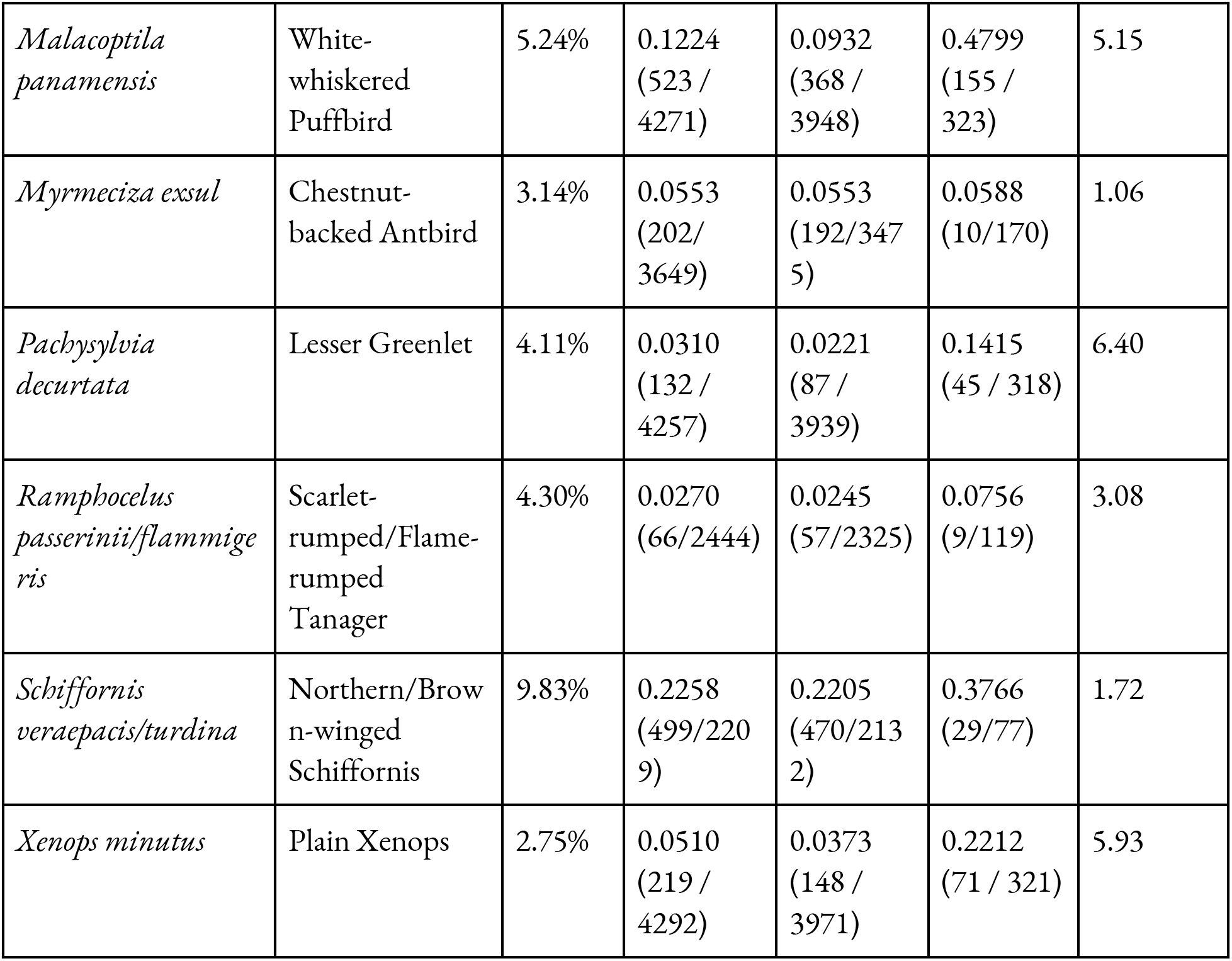
Proportion of fixed SNPs by taxon, broken down by autosomal and sex chromosomes. mtDNA distance calculated as pairwise difference of all coding regions.

| Taxon |  | mtDNA distance | Total proportion fixed SNPs | Proportion fixed autosomal SNPs | Proportion fixed Z-linked SNPs | Proportion Z vs autosomal fixed SNPs |
| --- | --- | --- | --- | --- | --- | --- |
| *Arremon aurantirostris* | Orange-billed Sparrow | 6.36% | 0.0202 (40/1979) | 0.0156 (29/1836) | 0.0887 (11/124) | 5.68 |
| *Cantorchilus nigricapillus* | Bay Wren | 3.30% | 0.0126 (37/2935) | 0.0105 (29/2757) | 0.0449 (8/178) | 4.28 |
| *Cyanocompsa cyanoides* | Blue-black Grosbeak | 5.01% | 0.0214 (85 / 3975) | 0.0172 (64 / 3718) | 0.0817 (21 / 257) | 4.75 |
| *Henicorbina leucosticta* | White-breasted Wood-wren | 5.94% | 0.0151 (28 / 1855) | 0.0119 (21 / 1763) | 0.0761 (7 / 92) | 6.39 |
| *Malacoptila panamensis* | White-whiskered Puffbird | 5.24% | 0.1224<br>(523 / 4271) | 0.0932<br>(368 / 3948) | 0.4799<br>(155 / 323) | 5.15 |
| *Myrmeciza exsul* | Chestnut-backed Antbird | 3.14% | 0.0553<br>(202/ 3649) | 0.0553<br>(192/347 5) | 0.0588<br>(10/170) | 1.06 |
| *Pachysylvia decurtata* | Lesser Greenlet | 4.11% | 0.0310<br>(132 / 4257) | 0.0221<br>(87 / 3939) | 0.1415<br>(45 / 318) | 6.40 |
| *Ramphocelus passerinii/flammigeris* | Scarlet-rumped/Flame-rumped Tanager | 4.30% | 0.0270<br>(66/2444) | 0.0245<br>(57/2325) | 0.0756<br>(9/119) | 3.08 |
| *Schiffornis veraepacis/turdina* | Northern/Brown-winged Schiffornis | 9.83% | 0.2258<br>(499/220 9) | 0.2205<br>(470/213 2) | 0.3766<br>(29/77) | 1.72 |
| *Xenops minutus* | Plain Xenops | 2.75% | 0.0510<br>(219 / 4292) | 0.0373<br>(148 / 3971) | 0.2212<br>(71 / 321) | 5.93 |

### Population structure and admixture detection

In all taxa, the Evanno method indicated that *K*=2 provided the best fit for our STRUCTURE results. Individuals with admixture proportions greater than 1% were detected in seven of the ten taxa, including all taxa considered conspecific across Panama except *Ma. panamensis* (Figure 2). When observed, the geographic extent of admixture varied widely (Figure 2) from admixed individuals occurring across our sampling transect in Panama, such as in *A. aurantiirostris*, occurring across nearly 350 km, to being confined to a roughly 5 km span between Cerro Azul and Cerro Jefe in central Panama, as seen in *H. leucosticta*.

**Figure 2:**
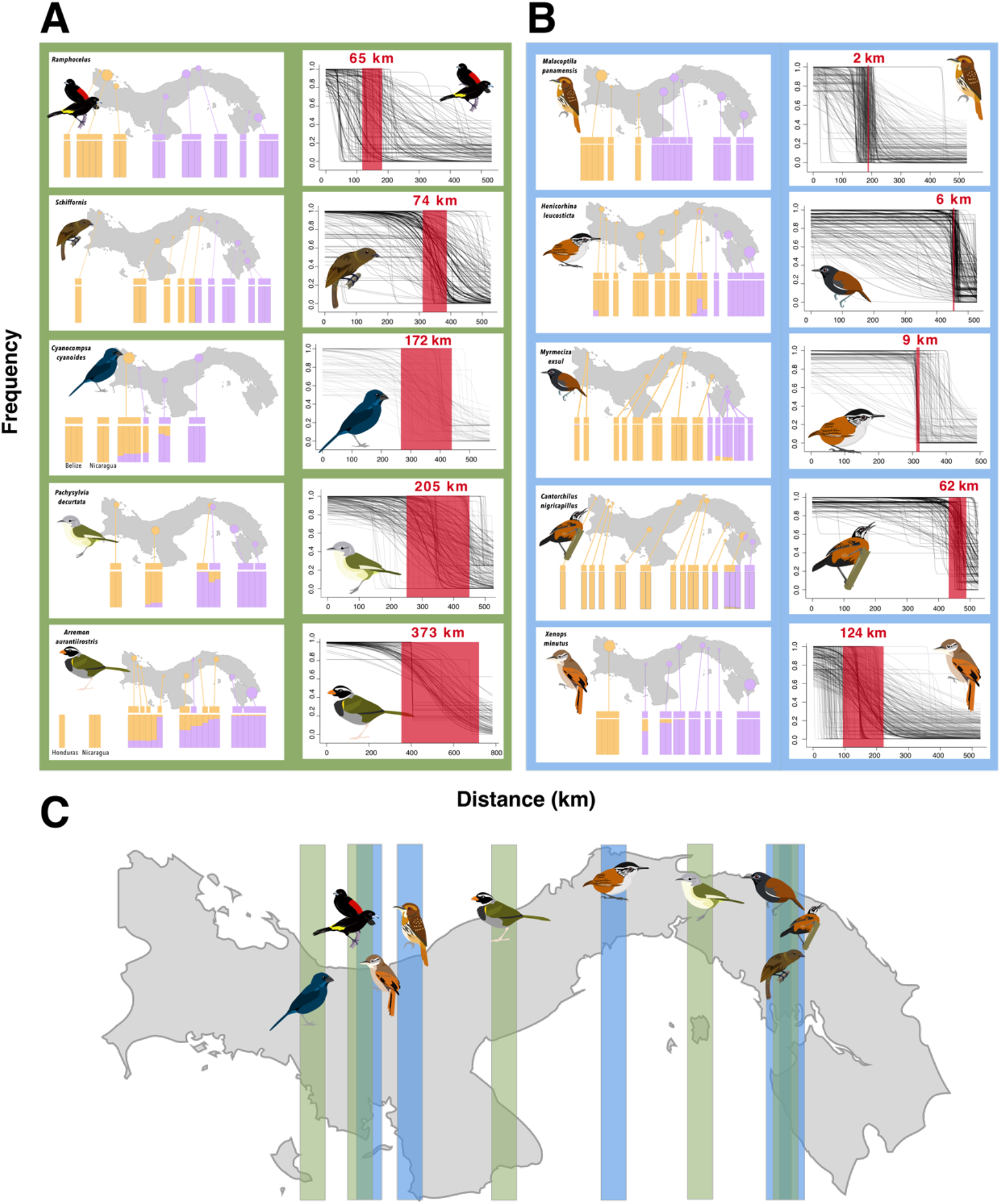
Genotypes of all birds, with points indicating locality, and cline results for the ten study taxa, group by diet guild with blue (A) indicating insectivores, and green (B) for granivores and frugivores. Structure plots in each taxon are grouped by locality, and bars on top indicate the mitochondrial haplotype of each individual. Genotypes for individuals from localities outside of Panama are labeled below each plot. Clines for each taxa are show per locus, with median clinal width highlighted in red. In C), approximate longitude of median cline centers for each taxa are mapped, with diet guild again indicated.

### Mitochondrial and nuclear mismatch

Mismatch between mitochondrial haplotype and nuclear genotype –defined as having the mitochondrial haplotype of one parental population with a nuclear genotype above 50% derived from the other parental population – was limited to three lineages (Figure 2). In *P. decurtata*, two individuals near the median cline center for that lineage had haplotypes characteristic of the parental population that was less than 50% of the nuclear genome. In *Ca. nigricapillus*, samples that had nuclear genotypes almost completely assignable to the Darién group nevertheless had the western mitochondrial haplotype. Again, this occurred very close to the median clinal center, and at one locality included only two of the three birds from the location. Finally, the central Panama localities for *A. aurantiirostris* were admixed with the greater assignment probability to the eastern Darien population, but with one exception, had the western haplotype.

### Cline analysis

Variation in geographic cline width and center provides information about the permeability of genomes to introgression. More permeable genomes are expected to show greater cline width and have a wider span of cline centers; narrow cline width and tightly coincident cline centers are a signature of RI (75–80). For both geographic and genomic cline analyses, our reduced diagnostic-only SNP datasets-- SNPs with a frequency of one genotype of 0.75 or more in one parental population and 0.25 or less in the other-- included between 72 (*A. aurantiirostris*) and 717 (*Ma. panamensis*) SNPs (Table 2). Median width varied from 2.4 km (*Ma. panamensis*) to 373.6 km (*A. aurantiirostris*; Table 2; Figure 2). Within each taxon, the widths best fit a Poisson distribution (Figure S1), with most cline widths very narrow and with a long tail of wider clines and the overall proportion of the loci in this tail varying across taxa. The median cline centers were likewise wide ranging (Figure 2C), but normally distributed (Figure S2). When distance in kilometers from the beginning of a taxon’s transect was converted back into degrees longitude, this estimate ranged between −78.22 and −81.63 degrees, corresponding approximately with the western edge of the Valiente Peninsula and the western edge of Darien province, respectively (Figure 2C). Within this area, two clusters of cline centers were observed. The first, including *Schiffornis*, *My. exsul*, and *Ca. nigricapillus*, occur near the border of Darien, with the three occurring very closely together. The other, less tightly clustered, group, occurs along the Caribbean coast of Veraguas, and includes *Ramphocelus*, *Ma. panamensis, X. minutus*, and *Cy. cyanoides*.

**Table 2:** Clinal characters of all taxa, including number of diagnostic loci for each set of analyses.

| Taxon | Diagnostic loci | Median Cline Width (km) | Variance in Cline Width | Median cline center (degrees longitude) | Variance in Cline Center |
| --- | --- | --- | --- | --- | --- |
| *Arremon aurantiirostris* | 72 | 373 | 43502 | -80.27414 | 6009 |
| *Cantorchilus nigricapillus* | 225 | 62.2 | 48930 | -78.22381 | 8955 |
| *Cyanocompsa cyanoides* | 81 | 172 | 52274 | -81.63481 | 5207 |
| *Henicorbhina leucosticta* | 163 | 9.3 | 19116 | -79.53446 | 4566 |
| *Malacoptila panamensis* | 717 | 2.4 | 14048 | -80.86800 | 2225 |
| *Myrmeciza exsul* | 389 | 6.1 | 36187 | -78.30600 | 7573 |
| *Pachysylvia decurtata* | 333 | 205 | 52330 | -78.98317 | 6285 |
| *Ramphocelus passerinii/flammigeris* | 162 | 65.3 | 28828 | -81.27781 | 8638 |
| *Schiffornis veraepacis/stenorhyncha* | 505 | 74.6 | 13165 | -78.30600 | 4254 |
| *Xenops minutus* | 509 | 124 | 32521 | -81.15118 | 5287 |

### Prediction testing

The best-fit models for our five response variables (median cline width, variance in single-locus clinal width, clinal center variance, autosomal fixation rate, and Z-chromosome fixation rate) varied. When we fit models for Z-chromosome and autosomal fixation rates, we found that mitochondrial distance and HWI were the best predictors in both cases (Table 3). For the primary cline variable-- the median width of clines across all loci-- diet alone was the best predictor, with weaker support for other models (Table 3). This was a weakly statistically supported relationship (*t*= −2.2662, *df* = 5.3811, *p* = 0.06904; Figure 3C), likely due to the overall small number of taxa included, but it is enough to say with reasonable certainty that insectivores have smaller median cline widths (mean = 40.887 km) than birds reliant on plant diets (mean = 178.35 km). For variance in clinal center (i.e., spatial coincidence of the centers of hybrid zones), HWI was the best predictor, although moderate support was also found for the following predictors, in order of increasing delta AICc: *F_ST_* alone, mitochondrial distance + HWI, mitochondrial distance alone, and *π* + HWI (Table 3). This relationship was moderately statistically supported (Adj *R2* = 0.2995, *p* = 0.05884; Figure 3A). For variance in clinal width (i.e, how much the spatial extent of hybridization varied), *F_ST_* alone was the best predictor, with moderate support also found for *π* + *F_ST_*. However, this was not significant, with weak support found for the linear relationship between the two (Adj *R2* = 0.0719, *p* = 0.3323; Figure 3B). While these results were weakly significantly significant, much of this is likely due to the small number of taxa included relative to the variation, and though not definitive, is still indicative of a biologically significant relationship between these variables that bears further investigation with more taxa.

**Table 3:**
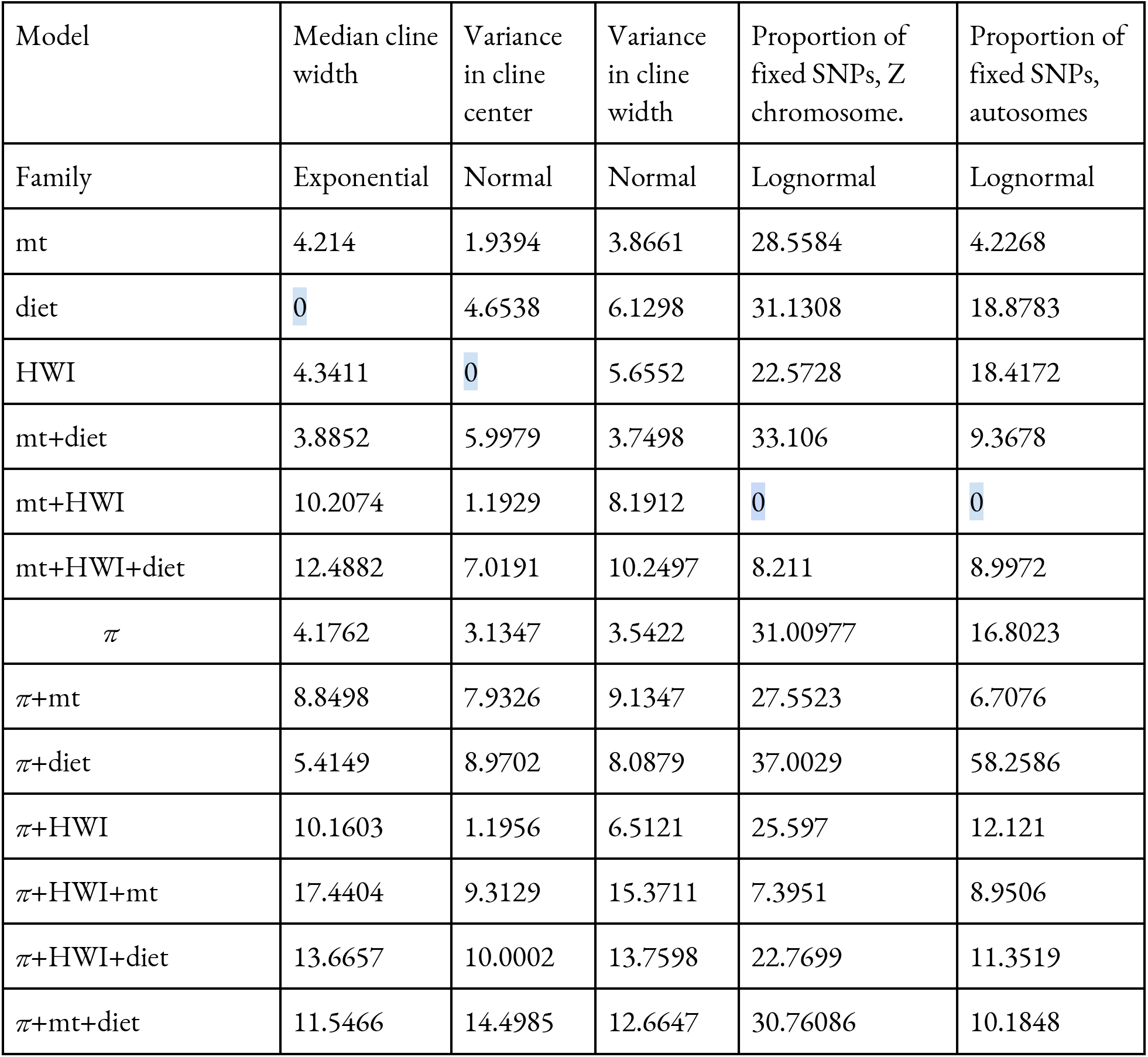
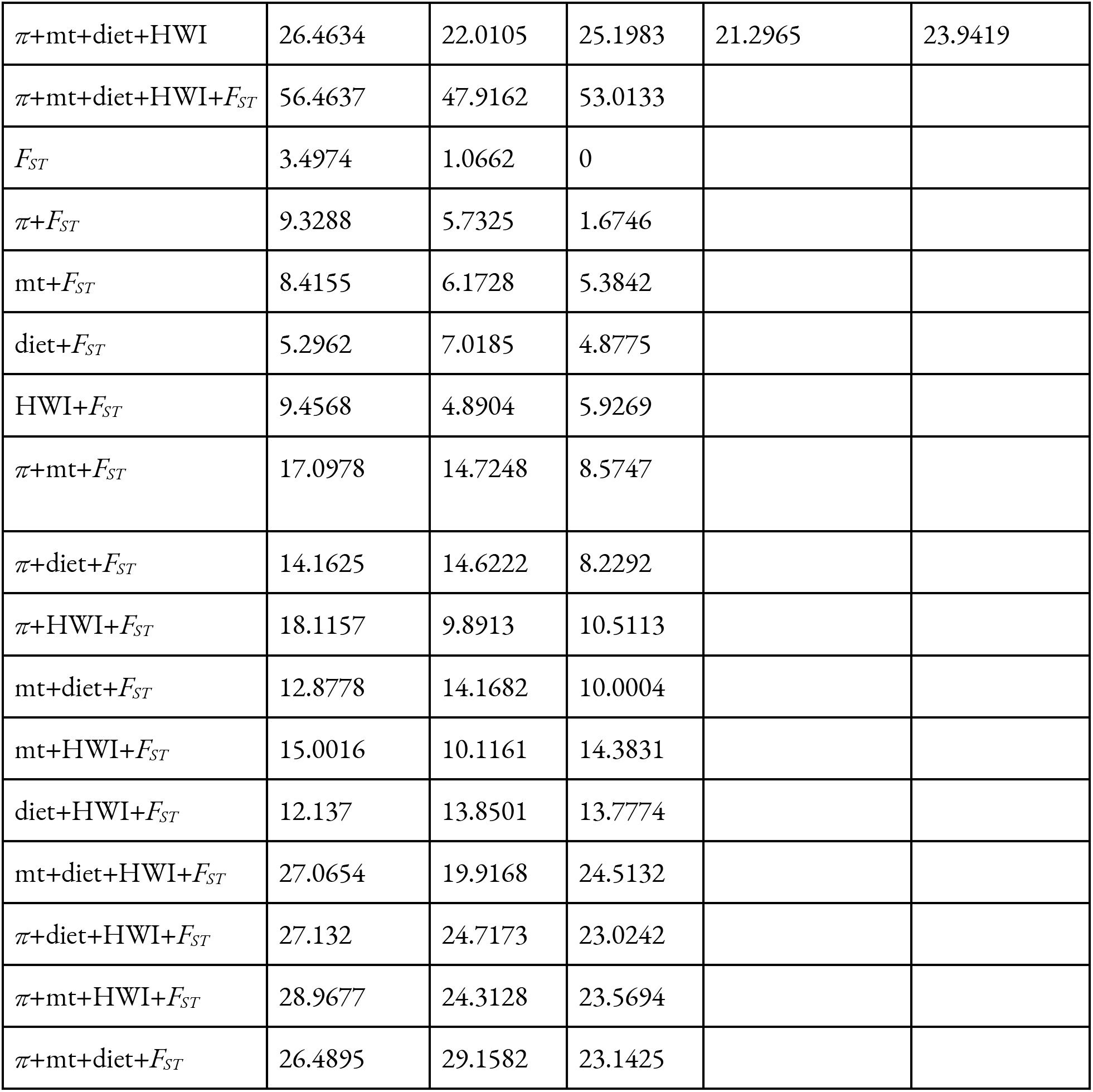
Delta AICc values for all tested GLMs. Model family for each response variable indicated.

| Model | Median cline width | Variance in cline center | Variance in cline width | Proportion of fixed SNPs, Z chromosome. | Proportion of fixed SNPs, autosomes |
| --- | --- | --- | --- | --- | --- |
| Family | Exponential | Normal | Normal | Lognormal | Lognormal |
| mt | 4.214 | 1.9394 | 3.8661 | 28.5584 | 4.2268 |
| diet | 0 | 4.6538 | 6.1298 | 31.1308 | 18.8783 |
| HWI | 4.3411 | 0 | 5.6552 | 22.5728 | 18.4172 |
| mt+diet | 3.8852 | 5.9979 | 3.7498 | 33.106 | 9.3678 |
| mt+HWI | 10.2074 | 1.1929 | 8.1912 | 0 | 0 |
| mt+HWI+diet | 12.4882 | 7.0191 | 10.2497 | 8.211 | 8.9972 |
| $\pi$ | 4.1762 | 3.1347 | 3.5422 | 31.00977 | 16.8023 |
| $\pi$+mt | 8.8498 | 7.9326 | 9.1347 | 27.5523 | 6.7076 |
| $\pi$+diet | 5.4149 | 8.9702 | 8.0879 | 37.0029 | 58.2586 |
| $\pi$+HWI | 10.1603 | 1.1956 | 6.5121 | 25.597 | 12.121 |
| $\pi$+HWI+mt | 17.4404 | 9.3129 | 15.3711 | 7.3951 | 8.9506 |
| $\pi$+HWI+diet | 13.6657 | 10.0002 | 13.7598 | 22.7699 | 11.3519 |
| $\pi$+mt+diet | 11.5466 | 14.4985 | 12.6647 | 30.76086 | 10.1848 |
| $\pi$+mt+diet+HWI | 26.4634 | 22.0105 | 25.1983 | 21.2965 | 23.9419 |
| $\pi$+mt+diet+HWI+$F_{ST}$ | 56.4637 | 47.9162 | 53.0133 | | |
| $F_{ST}$ | 3.4974 | 1.0662 | 0 | | |
| $\pi$+$F_{ST}$ | 9.3288 | 5.7325 | 1.6746 | | |
| mt+$F_{ST}$ | 8.4155 | 6.1728 | 5.3842 | | |
| diet+$F_{ST}$ | 5.2962 | 7.0185 | 4.8775 | | |
| HWI+$F_{ST}$ | 9.4568 | 4.8904 | 5.9269 | | |
| $\pi$+mt+$F_{ST}$ | 17.0978 | 14.7248 | 8.5747 | | |
| $\pi$+diet+$F_{ST}$ | 14.1625 | 14.6222 | 8.2292 | | |
| $\pi$+HWI+$F_{ST}$ | 18.1157 | 9.8913 | 10.5113 | | |
| mt+diet+$F_{ST}$ | 12.8778 | 14.1682 | 10.0004 | | |
| mt+HWI+$F_{ST}$ | 15.0016 | 10.1161 | 14.3831 | | |
| diet+HWI+$F_{ST}$ | 12.137 | 13.8501 | 13.7774 | | |
| mt+diet+HWI+$F_{ST}$ | 27.0654 | 19.9168 | 24.5132 | | |
| $\pi$+diet+HWI+$F_{ST}$ | 27.132 | 24.7173 | 23.0242 | | |
| $\pi$+mt+HWI+$F_{ST}$ | 28.9677 | 24.3128 | 23.5694 | | |
| $\pi$+mt+diet+$F_{ST}$ | 26.4895 | 29.1582 | 23.1425 | | |

**Figure 3:**
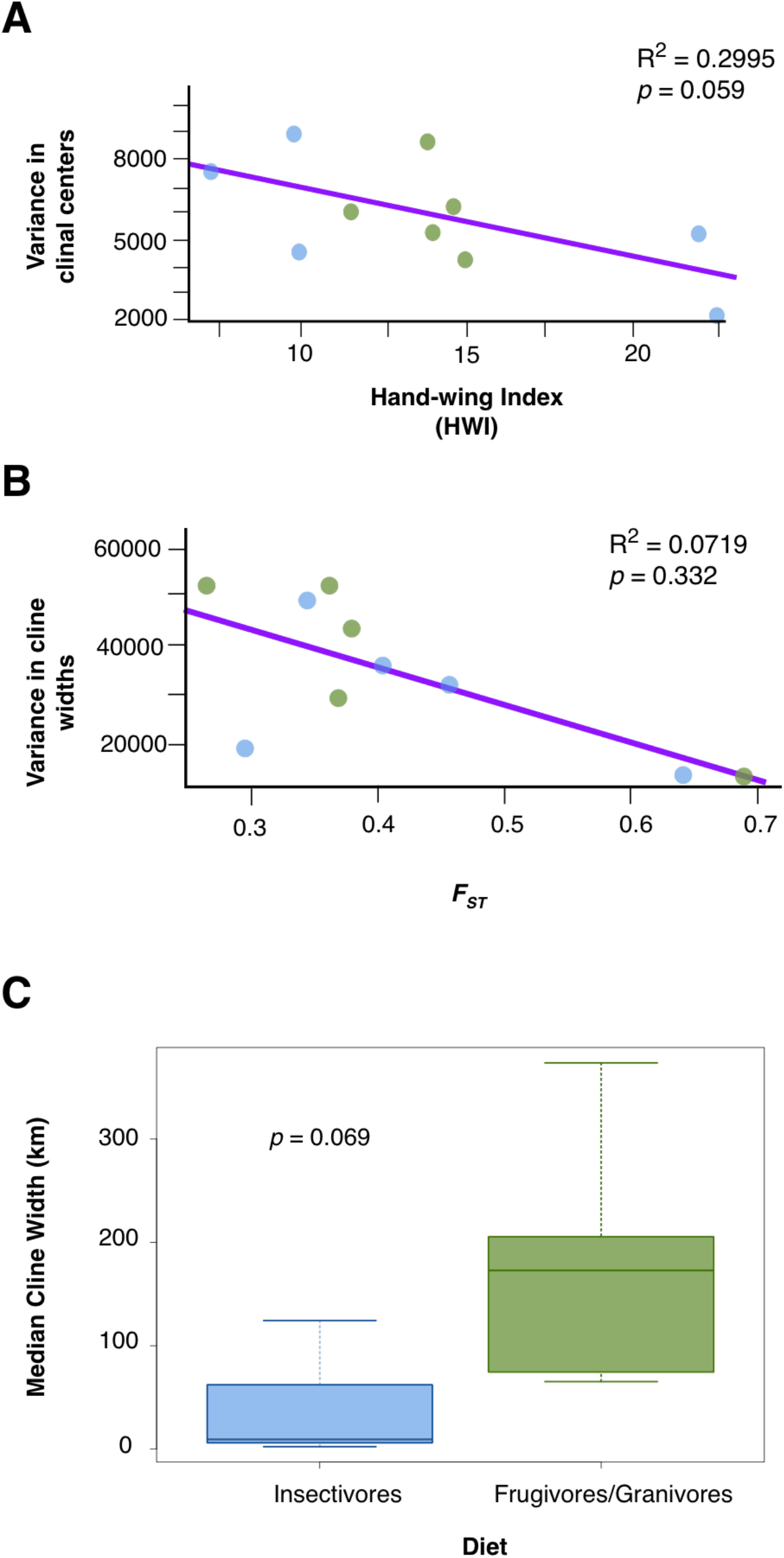
Best fit models for cline parameters. Plant-based diets indicated in green, insectivores in blue. A) For variance in cline center, HWI was the best fit, with a higher HWI corresponding with greater coincidence of the geographic center of clines (Adj *R^2^* = 0.2995, *p* = 0.05884). B) Variance in cline width was predicted instead by *F_ST_*, with cline widths becoming more consistent as overall genome differentiation increased, although this relationship was weak and not statistically supported (Adj *R^2^* = 0.0719, *p* = 0.3323). C) Median cline width was best predicted by diet type, with frugivores and granivores having wider clines than insectivores (*t*= −2.2662, *df* = 5.3811, *p* = 0.06904).

## Discussion

Time plays a role in predicting the accumulation of differentiation between populations. Yet time alone does not, in lowland Panama, tell the full story of how populations in secondary contact respond. Despite time often being implicitly assumed to play a key role in driving the development of RI in incipient species, our results show that time alone is not a predictor of these outcome of secondary contact in lowland Panama. It certainly plays a role in predicting the accumulation of fixed loci on both the autosomes and Z chromosome, supporting the mechanism of time allowing for the development of variation within isolated populations. Yet genomic differentiation alone does not necessarily indicate the development of RI, and for our most direct measure of the development of RI--median cline width-- time played no role as a predictor, indicating a disconnect between the mere accumulation of genomic variation and the development of differences that could lead to speciation.

### Predictors of geographic cline width and stability

We recovered diet alone as the best predictor of median cline width, placing ecological factors as key in determining the outcomes of secondary contact. Taxa that relied on plant-based foods, primarily seeds and/or fruit, had significantly wider clines than insectivores. Diet differences have been shown to be significantly linked with differences in dispersal ability in Neotropical birds (81, 82). Seeds and fruit are available year-round, but not necessarily in the same place simultaneously (83, 84). This results in birds reliant on such resources needing to disperse more widely (82). Meanwhile, insectivores, especially the understory insectivores examined here, can generally rely on a steady supply of food items year-round within a small area, and consequently are much more sedentary (85, 86) and have smaller home ranges and are often very weak fliers (81).

However, this link to dispersal ability is unlikely to be the sole reason for the strong relationship observed between diet and cline width. HWI, which reflects the morphological constraints on flight ability (88, 87) and more directly measures dispersal ability (85, 89–95), was not a predictor, so we must consider other demographic traits associated with diet that may be more influential in determining cline width. Notably, birds reliant on fruits and seeds have much wider variation in population sizes. Availability of these resources varies not just across a single year, but between years, with individual trees varying in fruit and seed production across multiple years (96), driving more pronounced cycles in population highs and lows in birds dependent on them (83, 97–100). While arthropods are also subject to declines, such cycles are generally relatively less drastic and usually specific to certain prey species (101). Because of this, Neotropical understory insectivores are observed to have far less year-to-year demographic fluctuation (98, 102–105). Thus, possibly the differences in demography among birds dependent on plant-based foods and those that primarily subsist on arthropods drive these patterns, although the limitations with UCE-linked loci make it difficult to test directly for historical demographic fluctuations.

Cline width alone is not the only indicator of the outcomes of secondary contact examined here. The coincidence of clines, as measured by the variation in cline center and width, is a measure of how spatially consistent the geographic center and extent of turnover will be across loci. Again, this was not predicted by time. The variance in cline center was best predicted by HWI alone. This is of interest because while dispersal ability as measured via the most accepted proxy was not important in determining the width of the hybrid zone, it is the best predictor behind how geographically concentrated introgression is. However, variance in width-- or how much variation in how far a locus introgresses across the transect-- was predicted by neither time nor ecological factors, but by *F_ST_*. This suggests that as overall genomic divergence between parental types increases, differential introgression of loci decreases, an observation corroborated by observations of decreased hybridization with greater genetic divergence in other systems (106).

The development of RI is a key step in the process of speciation, but it can be difficult to disentangle outside of controlled model systems, making hybrid zones an important opportunity to gain insight into the strength of RI in non-model organisms (22, 107, 108). Cline analyses can be powerful tools in gaining insight into how RI develops in the early stages of speciation (76, 79, 80). The geographic range of hybridization can be informative of the extent of RI in a pair of incipient species, as it reflects the extent to which introgressed loci are selected against (76, 79). Thus, median cline width across loci is a key measurement of the extent of RI.

### Geography of hybridization in lowland Panama

Of our ten taxa, seven of them had median cline centers in one of two narrow regions (Figure 2C). Three of the taxa-- *Schiffornis, My. exsul*, and *Ca. nigricapillus*-- had median cline centers estimated in western Darien, which also corresponded with where they experienced mitochondrial turnover. This corresponds to previous findings that this region represents a major suture zone between Central and South American taxa (109). The second cluster, consisting of the *Ramphocelus* tanagers, *Ma. panamensis, X. minutus*, and *Cy. cyanoides*, is located along the Caribbean coast of Veraguas. This region, whose avifauna has been much less well-documented than much of Panama (110–116), appears to be an important suture zone (117). In both cases, though, it is notable what they each lack: major biogeographic barriers that would easily explain the occurrence of such rapid turnover, a feature observed in other Neotropical lowland contact zones (118–123). Because of the apparent absence of barriers, these suture zones provide promising windows into how Neotropical diversity is shaped not by the landscape alone, but by traits inherent to ecological characteristics of the taxa in contact.

### Predictors of genomic differentiation

We found that time plays a role in generating variation across the genome, particularly the autosomes. This is consistent with predictions of neutral theory (124–126), as most of the genome accumulates variation at a predictable rate (127, 128). But just as there are fluctuations in the molecular clock that erode any simple link between the accumulation of mutations and time (128–130), the relationship between time and fixation rate is also less straightforward. While time was a significant predictor of autosomal fixation in our best-fit model, HWI was as well. The impact of time is clearly important-- in addition to being a predictor in our best-fit model, time alone was the second best-fit model for autosomal fixation rate (Table 3) -- but in this system, the relationship between the time and fixation is modulated by the dispersal capability. This is consistent with previous findings that high dispersal ability can hamper divergence in tropical birds (90, 95) but shows that this also extends to the accumulation of neutral variation.

The Z-chromosome is often the focus of searches for loci driving the development of RI (49, 65, 66, 69, 131). The accumulation of fixed differences on the sex chromosomes has implications for the development of RI (49, 65, 132, 133), as they can lead to incompatibilities (49, 134) or be linked to traits involved in male-biased (in ZW systems) sexual selection or conflict (135–137). We found that while time was a predictor of Z-chromosome differentiation, it was not significant, while the other included predictor, HWI, was. Dispersal ability, of which HWI is a well-established proxy (85, 90, 94, 95), then, appears to play a role in driving divergence in the specific portion of the genome most frequently implicated in speciation in birds (49, 65, 67). As dispersal ability has been previously shown to play a major role in shaping hybrid zone dynamics (138), this is not unexpected. However, it is notable that this was the only response variable for which dispersal ability itself, as measured with HWI, is a predictor.

### Time drives the generation of variation, but not of RI

Our results confirm that time does play a role in the development of genomic differentiation, those differences do not necessarily indicate the development of RI. Disentangling how genomic variation, phenotypic differences, and reproductive barriers are intertwined can prove challenging. Previous work in birds, including wood-warblers (139), hummingbirds (26), and seedeaters (140) show that remarkably little genomic variation can nevertheless result in striking phenotypic variation. Yet most such examples recover evidence of substantial gene flow that indicate that such differences should not in and of themselves be taken as evidence for RI, which has been backed up in other systems where overall genetic differentiation has not necessarily been indicative of the extent of reproductive barriers (141, 142; but see 143 for counterexample).

### Lessons from secondary contact on the formation of species

Hybrid zones have long been acknowledged as a powerful window into the evolutionary mechanisms driving speciation (107, 108, 144–146). Regions like lowland Panama, where hybrid zones in multiple taxa co-occur, have even greater potential for investigation of broader questions about the speciation process. However, many such studies tend to focus on questions of how landscape factors across time have impacted divergence in large suites of organisms (27). Lowland Panama provides an important setting for a different set of questions. As the range of split times suggests no single historic factor and the region lacks the strong geographic barriers characteristic of other classic suture zones, the spotlight can instead be shifted onto the inherent traits of the organisms themselves. Our results drive home the necessity of considering ecological factors such as demographic stability and dispersal capability as key drivers of outcomes of contact between diverging populations.

## Materials and methods

### Taxonomic and genetic sampling

The avifauna of the lowlands of Panama provide an excellent system to test hypotheses about how time shapes the outcomes of secondary contact. Connected by largely continuous forest prior to the relatively recent arrival of settler-colonial agricultural practices (147, 148) and lacking obvious north-south geographic barriers between the eastern and western lowlands (121), this region is nevertheless a notable suture zone where many sister taxa rapidly replace each other, with varying degrees of apparent hybridization (149). We focused on taxa that are found widely across this region and for which genetic resources from vouchered museum specimens were available (Table S1). We also considered preliminary data from mitochondrial barcoding (149) to include as wide a range of split depths as possible, along with diet, habitat, and family, to include an ecologically diverse sample of species. This resulted in a final dataset of ten lineages with mtDNA divergence between 2.75% and 9.83% (pairwise distance of all mt coding regions).

We examined loci associated with UCEs to measure clinal variation in secondary contact in hundreds or thousands of homologous loci, and to estimate the degree of genome-wide fixed variation between populations at terminal populations across transects. UCEs are useful for comparative studies, as they allow for robust sampling of directly comparable orthologous loci across large taxonomic divides (27, 150–152). For each taxon, we attempted to sample 3-4 individuals per population from populations along an east-west transect across lowland Panama, but in several cases were only able to include one or two individuals from a locality, resulting in a dataset of 180 individuals (Table S1). Preliminary analyses suggested that two species— *Cyanocompsa cyanoides* and *Arremon aurantiirostris*—had signals of admixture in the westernmost population sampled in northwest Panama. Therefore, for these species we sampled added populations from Nicaragua and Honduras for both species, and Belize for *Cy. cyanoides*. Similarly, we added samples to the east from coastal Ecuador for *Cantorchilus nigricapillus*, ensuring that all 10 species had purely parental populations in the westernmost and easternmost terminal populations.

We extracted genomic DNA from muscle tissue samples from vouchered museum specimens (Table S1) and prepared UCE-enriched libraries following Faircloth *et al*. (152) and Glenn *et al*. (153). For a few individuals, UCEs were harvested from whole-genome shotgun sequenced libraries. In both cases, libraries were prepared following the NEB Ultra II protocol. For the UCE-enriched libraries, we then followed the protocol from Glenn *et al* (153) using the 5k Tetrapod set v.1 of 5,060 probes (152), while non-enriched samples proceeded directly to sequencing (Table S1).

### Bioinformatics

We used either Trimmomatic v. 0.32 (154; implemented in Illumiprocessor, 155) or BBDuk (part of BBMap v. 37.93; 156) for quality checking and adapter-trimming in raw reads, with the former being used for UCE-only reads as part of the original UCE pipeline and the latter for our whole genome reads. We generated a UCE loci-only reference for each taxon with *de novo* assemblies using Trinity v. 2.5.1 (157), filtering the assembly contigs to ensure that they contained only single-copy UCE loci without capture bycatch with phyluce (158). Read number is not necessarily predictive of enrichment quality, so we generated a UCE assembly for all individuals, and selected the assembly with the most recovered UCE loci as the reference for that taxon. However, for three taxa, we had no UCE-enriched libraries. Assuming that the number of total reads was proportional to UCE-derived reads in these samples, we selected the single individual with the greatest number of reads to generate a reference for that taxon. For that sample’s trimmed reads, we filtered the UCE-derived reads using BBSplit (part of BBMap) and a concatenated reference derived from a set of closely related taxa as the BBSplit mapping target. For *Myrmeciza exsul* and *Schiffornis*, the three closely related taxa were *Xenops minutus, Henicorhina leucosticta*, and *Mionectes oleagineus* (developed for separate project); for *Ramphocelus*, we used *Pachysylvia decurtata*, *Cy. cyanoides*, and *A. aurantiirostris*. Using the BBSplit reads, we generated a final reference pseudo-genome for each species, as above.

To recover diploid genotypes for all UCE loci, we followed the Genome Analysis Toolkit (GATK; 159) best practices, making an alignment using all reads mapped to the reference using BWA (160, 161). The alignment was indexed and cleaned using Samtools v. 1.7 (162) and Picard v. 2.18.0 (163). Genotypes were generated in GATK v. 4.0.12 using HaplotypeCaller and GenotypeGVCF, generating a variant file per taxon. Using VCFTools v. 0.1.13 (164), the variants were filtered to include only variants with biallelic SNPs present in all individuals with a minimum GQ of 10 and mean coverage depth of 10. This filtered dataset was used for window-based analyses of nucleotide diversity and genome-wide *F_ST_*. We further thinned this dataset to include one SNP per locus by using the thin function of VCFTools, retaining the first 5’ SNP for each locus. This became our canonical SNP dataset for each taxon for non-windowed analyses. We also identified Z-linked loci to test how fixation rates differed between autosomal and sex-linked chromosomes. The master UCE probe sequence list (152) was matched to the *Taeniopygia guttata* genome (GCA_008822105.2) with BLAST (165) to generate a list of UCE loci located on the Z chromosome, as chromosomal synteny in birds is relatively conserved (49, 56, 55). This list was then used to reference against per-site *F_ST_ to* determine autosomal versus Z-linked fixation.

#### Determining the mitochondrial haplotype

We assembled mitochondrial genomes for both the eastern and western parental populations in each with NOVOplasty v. 3.4 (166). We then determined haplotype per individual (149).

### Population structure analyses

To verify that each species included just two parental genotypes across the study area, to ensure that each parental genotype was adequately sampled, and to provide an initial assessment of population structure, for each species we conducted PCA and DAPC in the R package adegenet v. 2.1.3 (167). We determined the number of clusters by minimizing the value of the Bayesian Information Criterion (BIC), and individuals were assigned to these clusters. We used STRUCTURE v2.3.4 (168) to provide an additional assessment of population structure, and to assign individuals as either the western or eastern parental genotypes or as admixed between the two. We ran STRUCTURE for 30 replicates for each of *K* = 2–4. We combined all replicates for each *K* using CLUMPP v. 1.12 (169). We confirmed the best value of *K* as 2 in STRUCTURE HARVESTER v. 0.6.94 (170) using the Evanno method (171), and plotted results with distruct (172).

### Geographic cline width analysis

We generated geographic clines as follows: we filtered our SNP datasets to include only diagnostic loci (48), defined as loci where the westernmost populations had a combined allele frequency for genotype *p* of at least 0.75, while the combined allele frequency in the easternmost populations was no more than 0.25 for *p*. We generated geographic clines for these diagnostic loci using the R package hzar (76), testing 3 cline models per locus using a custom script (https://github.com/jfmclaughlin92/panama_uce). Using the best-fit model for each locus, we calculated median cline center and width for each taxon, as well as the set of cline width and center values that represent 95% of the observed variation. We then plotted all best-fit clines together for each taxon, calculating the median width and the variance of the width and center across all loci.

#### Statistics and prediction testing

We obtained counts of fixed SNPs using VCFTOOLS, including across all loci, autosomal sites only, and Z-linked loci only, as determined by alignment of the UCE probe sequences to the zebra finch genome. We then constructed a general linear model for each response parameter. In addition to mitochondrial distance, we also included variables to test for how other factors contribute to shaping the outcomes of secondary contact. We included hand-wing index (HWI), a quantification of the aspect ratio of a bird’s wing that is an indicator of dispersal ability, a known driver of divergence in Neotropical birds (90). We additionally included diet, coded as plant-based (frugivores and granivores) or insectivorous. These diet types are linked to differing dispersal capabilities (121) and characteristic demographic patterns (82, 84, 173, 174). Finally, weighted genome-wide *F_ST_* was included as a generalized measurement of genomic differentiation, and *π* for overall nucleotide diversity. These were calculated from only the parental populations, and clines were not calculated from them in such a way to make them autocorrelated. For cline parameters (median width, variance in width, and variance in center), these were *F_ST_*, mitochondrial distance, HWI (94), and diet. For autosomal and Z-linked fixation rates, we excluded *F_ST_*, as that would be auto-correlated with the response variables. We tested which error structure best described each dataset, and from there constructed generalized linear models (GLMs) using all combinations of the explanatory variables. From these models produced for each response variable, we then used AICc to select the best model.

## Acknowledgements

This project was supported by the Ornithology Department of the Sam Noble Museum and the Sutton Scholarship Fund at the University of Oklahoma. Scientific collecting permits for birds collected for this study were obtained from MiAmbiente, who we thank for their support of scientific collecting. We thank the University of Alaska Museum, University of Washington Burke Museum, and Smithsonian Tropical Research Institute Bird Collection for providing tissue grants from museum specimens used in this study. We would like to thank Peggy Guitton-Mayerma and Jorge Luis Garzon for field work; Celestino Aguilar, Graham Wiley, and Kevin Hawkins for assisting in lab work; Sara Lipshutz, for generously sharing scripts for running hzar; and Dahiana Arcila, Hayley Lanier, Courtney Hofman, JP Masly, Kaiya Provost, Paige Byerly, Auriel Fournier, Anusha Bishop, and Valentina Gomez for their feedback on earlier versions of the manuscript.

## Notes

### Competing Interest Statement

The authors have declared no competing interest.

## References

1. E. Mayr. Systematics and the Origin of Species, from the Viewpoint of a Zoologist. (Harvard University Press, 1942).

2. E. Mayr. Animal Species and Evolution. (Harvard University Press, 1963).

3. G. L. Bush. Modes of animal speciation. Annu. Rev. Eco. Syst., 6, 339–364 (1975).

4. J. D. Lynch. “The gauge of speciation: on the frequency of modes of speciation” in Speciation and its consequences, D. Otte, J. A. Endler, Eds. (Sinauer Associates Inc, 1989). pp []

5. D. Otte, J. A. Endler. Speciation and its consequences. Sinauer Associates Inc, Sunderland, MA, USA. (1989).

6. M. L. Rosenzweig. Species diversity in space and time. (Cambridge University Press, 1995).

7. T. G. Barraclough, A. P. Vogler. Detecting the geographical pattern of speciation from species-level phylogenies. Am. Nat., 155, 419–434 (2000).

8. J. A. Coyne, H. A. Orr. Speciation. (Sinauer Associates, Inc, 2004).

9. C. I. Wu The genic view of the process of speciation. J. Evol. Biol., 14, 851–865 (2001).

10. J. Hey, R. S. Waples, M. L. Arnold., R. K. Butlin, R. G. Harrison. Understanding and confronting species uncertainty in biology and conservation. Trends Ecol. Evol., 18, 597–603 (2003).

11. J. Mallet, M. Beltrán, W. Neukirchen, M. Linares, Natural hybridization in Heliconiine butterflies: the species boundary as a continuum. BMC Evol. Biol., 7, 28 (2007).

12. P. Nosil, D. J. Funk, D. Ortiz-Barrientos. Divergent selection and heterogeneous genomic divergence. Mol. Ecol., 18, 375–402 (2009).

13. O. Seehausen, et al. Genomics and the origin of species. Nat. Rev. Genet., 15, 176–192 (2014).

14. W. C. Webb, J. M. Marzluff, K. E. Omland. Random interbreeding between cryptic lineages of the common raven: evidence for speciation in reverse. Mol. Ecol., 20, 2390–2402 (2011).

15. P. R. Grant, B. R. Grant. Introgressive hybridization and natural selection in Darwin’s finches. Biol. J. Linn. Soc., 117, 812–822 (2016).

16. A. M. Kearns, et al. Genomic evidence of speciation reversal in ravens. Nat. Comm., 9, 906 (2018).

17. D. L. Slager, et al. Cryptic and extensive hybridization between ancient lineages of American crows. Mol. Ecol., 29, 956–969 (2020).

18. S. A. Cowles, J. A. C Uy. Rapid, complete reproductive isolation in two closely related *Zosterops* white-eye bird species despite broadly overlapping ranges. Evolution, 73, 1647–1662 (2019).

19. C. Merot, C. Salazar, R. M. Merrill, C. D. Jiggins, M. Joron. What shapes the continuum of reproductive isolation? Lessons from Heliconius butterflies. Proc. Roy. Soc. B, 284, 20170335 (2017).

20. A. P. Hendry, D. I. Bolnick, D. Berner, C. L. Peichel. Along the speciation continuum in sticklebacks. J. Fish Biol., 75, 2000–2036 (2009).

21. J. Mallet. Hybridization as an invasion of the genome. Trends Ecol. Evol., 20, 229–237 (2005).

22. J. Mallet, K. K. Dasmahapatra. Hybrid zones and the speciation continuum in *Heliconius* butterflies. Mol. Ecol., 21, 5643–5645 (2012).

23. P. Nosil. Speciation with gene flow could be common. Mol. Ecol., 17, 2103–2106 (2008).

24. S. H. Martin, et al. Genome-wide evidence for speciation with gene flow in *Heliconius* butterflies. Genome Res., 23, 1817–1828 (2013).

25. J. Ottenburghs, et al. A history of hybrids? Genomic patterns of introgression in the true geese. BMC Evol. Biol., 17, 201 (2017).

26. C. Palacios et al. Shallow genetic divergence and distinct phenotypic differences between two Andean hummingbirds: speciation with gene flow? Auk, 136, ukz046 (2019).

27. J. F. McLaughlin, B. C. Faircloth, T. C. Glenn, K. Winker. Divergence, gene flow, and speciation in eight lineages of trans-Beringian birds. Mol. Ecol., 29, 3526–3542 (2020).

28. H. A. Orr. The population genetics of speciation: the evolution of hybrid incompatibilities. Genetics, 139, 1805–1813 (1995).

29. S. Singhal, C. Moritz. Reproductive isolation between phylogeographic lineages scales with divergence. Proc. Roy. Soc. B, 280, 20132246 (2013).

30. H. A. Orr, M. Turelli. The evolution of postzygotic isolation: accumulating Dobzhansky-Muller incompatibilities. Evolution, 55, 1085–1094 (2001).

31. G. L. Bush, S. M. Case, A. C. Wilson, J. L. Patton. Rapid speciation and chromosomal evolution in mammals. Proc. Natl. Acad. Sci., 74, 3942–3946 (1977).

32. E. L. Berdan, A. Blanckaert, R. K. Butlin, C. Bank. Deleterious mutation accumulation and the long-term fate of chromosomal inversions. biorXiv [Preprint] (2020). https://doi.org/10.1101/606012 (Accessed 17 November 2021).

33. G. E. Hill. Mitonuclear coevolution as the genesis of speciation and the mitochondrial DNA barcode gap. Ecol. Evol., 6, 5831–5842 (2016).

34. G. E. Hill. The mitonuclear compatibility species concept. Auk, 134, 393–410 (2017).

35. S. Wright. Evolution in Mendelian populations. Genetics, 16, 97–159 (1931).

36. M. M. Sasa, P. T. Chippindale, N. A. Johnson. Patterns of postzygotic isolation in frogs. Evolution, 52, 1811–1820 (1998).

37. P. L. Tubaro, D. A. Lijtmaer Hybridization patterns and the evolution of reproductive isolation in ducks. Biol. J. Linn. Soc., 77, 193–200 (2002).

38. D. A. Lijtmaer, B. Mahler, P. L. Tubaro. Hybridization and postzygotic isolation patterns in pigeons and doves. Evolution, 57, 1411–1418 (2003).

39. D. I. Bolnick, T. J. Near. Tempo of hybrid inviability in centrarchid fishes (Teleostei: Centrarchidae). Evolution, 59, 1754–1767 (2005).

40. C. Dufresnes et al. Timeframe of speciation inferred from secondary contact zones in the European tree frog radiation (*Hyla arborea* group). BMC Evol. Biol., 15, 155 (2015).

41. T. D. Price Speciation in Birds. (Roberts and Company, 2008).

42. T. D. Price, M. M. Bouvier. The evolution of F1 postzygotic incompatibilities in birds. Evolution, 56, 2083–2089 (2002).

43. B. M. Winger. Consequences of divergence and introgression for speciation in Andean cloud forest birds. Evolution, 71, 1815–1831 (2017).

44. B. M. Winger, J. M. Bates. The tempo of trait divergence in geographic isolation: avian speciation across the Marañon Valley of Peru. Evolution, 69, 772–787 (2015).

45. S. V. Edwards et al. (2005). Speciation in birds: genes, geography, and sexual selection. Proc. Natl. Acad. Sci., 102, 6550–6557

46. B. M. Fitzpatrick. Rates of evolution of hybrid inviability in birds and mammals. Evolution, 58, 1865–1870 (2004).

47. J. M. Gee. Gene flow across a climatic barrier between hybridizing avian species, California and Gambel’s quail (*Callipepla californica* and *C. gambelii*).” Evolution, 58, 1108–1121 (2004).

48. S. E. Lipshutz. et al. Differential introgression of a female competitive trait in a hybrid zone between sex-role reversed species. Evolution, 73, 188–201 (2019).

49. H. Ellegren. The evolutionary genomics of birds. Annu. Rev. Ecol. Evol S., 44, 239–259 (2013).

50. K. Bachmann, B. A. Harrington, J. P. Craig. Genome size in birds. Chromosoma, 37, 405–416 (1972).

51. T. R. Tiersch, S. S. Wachtel. On the evolution of genome size of birds. J. Hered., 82, 363–368 (1991).

52. N. A. Wright, T. R. Gregory, C. C. Witt. Metabolic ‘engines’ of flight drive genome size reduction in birds. Proc. Roy. Soc. B, 281, 20132780 (2014).

53. T. R. Gregory. A bird’s-eye view of the C-value enigma: genome size, cell size, and metabolic rate in the class Aves. Evolution, 56, 121–130 (2002).

54. A. Kapusta, A. Suh, C. Feschotte. Dynamics of genome size evolution in birds and mammals. Proc. Natl. Acad. Sci., 114, 1460–1469 (2017).

55. S. Shetty, D. K. Griffin, Graves, J. A. Comparative painting reveals strong chromosome homology over 80 million years of bird evolution. Chromosome Res., 7, 289–295 (1999).

56. H. Ellegren. Evolutionary stasis: the stable chromosomes of birds. Trends Ecol. Evol., 25, 283–291 (2010).

57. D. E. Irwin. Assortative mating in hybrid zones is remarkably ineffective in promoting speciation. Am. Nat., 195, E150–167 (2020).

58. E. Nemeth et al. Bird song and anthropogenic noise: vocal constraints may explain why birds sing higher-frequency songs in cities. Proc. Roy. Soc. B, 280, 20122798 (2013).

59. T. J. Parsons, S. L. Olson, M. J. Braun. Unidirectional spread of secondary sexual plumage traits across an avian hybrid zone. Science, 260, 1643–1646 (1993).

60. R. T. Brumfield, M. J. Braun. Phylogenetic relationships in bearded manakins (Pipridae: *Manacus*) indicate that male plumage color is a misleading taxonomic marker. Condor, 103, 248 (2001).

61. T. L. Parchman et al. The genomic consequences of adaptive divergence and reproductive isolation between species of manakins. Mol. Ecol., 22, 3304–3317 (2013).

62. A. C. Stein, J. A. C. Uy. Unidirectional introgression of a sexually selected trait across an avian hybrid zone: a role for female choice? Evolution, 60, 1476–1485 (2006).

63. D. T. Baldassarre, M. S. Webster. Experimental evidence that extra-pair mating drives asymmetrical introgression of a sexual trait. Proc. Roy. Soc. B, 280, 20132175 (2013).

64. D. T. Baldassarre, T. A. White, J. Karubian, M. S. Webster. Genomic and morphological analysis of a semipermeable avian hybrid zone suggests asymmetrical introgression of a sexual signal. Evolution, 68, 2644–2657 (2014).

65. D. E. Irwin. Sex chromosomes and speciation in birds and other ZW systems. Mol. Ecol., 27, 3831–3851 (2018).

66. N. Backström et al. A high-density scan of the Z chromosome in *Ficedula* flycatchers reveals candidate loci for diversifying selection. Evolution, 64, 3461–3475 (2010).

67. P. Lavretsky et al. Speciation genomics and a role for the Z chromosome in the early stages of divergence between Mexican ducks and mallards. Mol. Ecol., 24, 5364–5378 (2015).

68. P. Lavretsky, J. M. Dacosta, M. D. Sorenson, K. G. McCracken, J. L. Peters. ddRAD-Seq data reveal significant genome-wide population structure and divergent genomic regions that distinguish the mallard and close relatives in North America. Mol. Ecol., 28, 2594–2609 (2019).

69. R. Storchová, J. Reif, M. W. Nachman. Female heterogamety and speciation: reduced introgression of the Z chromosome between two species of nightingales. Evolution, 64, 456–471 (2010).

70. J. B. S. Haldane. Sex ratio and unisexual sterility in hybrid animals. J. Genet., 12, 101–109 (1922).

71. C. C. Laurie. The weaker sex is heterogametic: 75 years of Haldane’s rule. Genetics, 147, 937–951 (1997).

72. J. A. Coyne. The genetic basis of Haldane’s Rule. Nature, 314, 736–738 (1985).

73. N. A. Johnson, J. Lachance. The genetics of sex chromosomes: evolution and implications for hybrid incompatibility. Ann. NY Acad. Sci., 1256, 1–22 (2012).

74. C. N. Trier, J. S. Hermansen, G. P. Sætre, R. I. Bailey. Evidence for mito-nuclear and sex-linked reproductive barriers between the hybrid Italian sparrow and its parent species. PLoS Genet., 10, e1004075 (2014).

75. N. H. Barton. Gene flow past a cline. Heredity, 43, 333–339 (1979).

76. E. P. Derryberry, G. E. Derryberry, J. M. Maley, R. T. Brumfield. HZAR: hybrid zone analysis using an R software package. Mol. Ecol. Resour., 14, 652–663 (2014).

77. R. G. Harrison. Hybrid zones: windows on evolutionary process. Oxford Surveys in Evolutionary Biology, 7, 69–128 (1990).

78. L. H. Rieseberg, J. Whitton, K. Gardner. Hybrid zones and the genetic architecture of a barrier to gene flow between two sunflower species. Genetics, 152, 713–727 (1999).

79. J. M. Szymura, N. H. Barton. Genetic analysis of a hybrid zone between the fire-bellied toads, *Bombina bombina* and *B. variegata*, near Cracow in southern Poland. Evolution, 40, 1141–1159 (1986).

80. J. M. Szymura, N. H. Barton. The genetic structure of the hybrid zone between the fire-bellied toads *Bombina bombina* and *B. variegata*: comparisons between transects and between loci. Evolution, 45, 237–261 (1991).

81. R. P. Moore, W. D. Robinson, I. J. Lovette, T. R. Robinson. Experimental evidence for extreme dispersal limitation in tropical forest birds. Ecol. Lett., 11, 960–968 (2008).

82. D. A. Westcott, D. L. Graham. Patterns of movement and seed dispersal of a tropical frugivore. Oecologia, 122, 249–257 (2000).

83. D. J. Levey, Moermond, T. C., Denslow, J. S. Frugivory: an overview. In: La Selva: Ecology and Natural History of a Neotropical Rain Forest. L. A. McDade, K. S. Bawa, H. A. Hespenheide, G. A. Hartshorn, Eds. (University of Chicago Press, 1994), pp. 282–294.

84. E. S. Morton. On the evolutionary advantages and disadvantages of fruit eating in tropical birds. Am. Nat., 107, 8–22 (1973).

85. C. W. Burney, R. T. Brumfield. Ecology predicts levels of genetic differentiation in Neotropical birds. Am. Nat., 174, 358–368 (2009).

86. D. J. Levey, F. G. Stiles. Evolutionary precursors of long-distance migration: resource availability and movement patterns in Neotropical landbirds. Am. Nat., 140, 447–476 (1992).

87. F. A. Kipp. Der handflügel-index als flugbiologisches maß. Die Vogelwarte, 20, 77–86 (1959).

88. R. Lockwood, J. P. Swaddle, J. M. V. Rayner. Avian wingtip shape reconsidered: wingtip shape indices and morphological adaptations to migration. J. Avian Biol., 29, 273–292 (1998).

89. V. L. Chua et al. Evolutionary and ecological forces influencing population diversification in Bornean montane passerines. Mol. Phylogenet. Evol., 113, 139–149 (2017).

90. S. Claramunt, E. P. Derryberry, J. V. Remsen, R. T. Brumfield High dispersal ability inhibits speciation in a continental radiation of passerine birds. Proc. Roy. Soc. B, 279, 1567–1574 (2012).

91. S. Claramunt, N. A. Wright. “Using museum specimens to study flight and dispersal” in The Extended Specimen: Emerging Frontiers in Collections-Based Ornithological Research, M. S. Webster, Ed. (CRC Press, 2017), pp. 127–141.

92. J. D. Kennedy et al. The influence of wing morphology upon the dispersal, geographical distributions and diversification of the Corvides (Aves; Passeriformes). Proc. Roy. Soc. B, 283, 20161922 (2016).

93. A. L. Pigot, W. Jetz, C. Sheard, J. A. Tobias. The macroecological dynamics of species coexistence in birds. Nat. Ecol. Evol., 2, 1112–1119 (2018).

94. C. Sheard et al. Ecological drivers of global gradients in avian dispersal inferred from wing morphology. Nat. Comm., 11, 2463 (2020).

95. B. C. Weeks, S. Claramunt. Dispersal has inhibited avian diversification in Australasian archipelagoes. Proc. Roy. Soc. B, 281, 20141257 (2014).

96. N. T. Wheelwright. “A seven-year study of individual variation in fruit production in tropical bird-dispersed tree species in the family Lauraceae” in Frugivores and Seed Dispersal, A. Estrada, T. H. Fleming, Eds. (Springer Netherlands, 1986), pp. 19–35

97. J. D. Brawn, J. R. Karr, J. D. Nichols. Demography of birds in a Neotropical forest: effects of allometry, taxonomy, and ecology. Ecology, 76, 41–51 (1995).

98. J. Faaborg, W. J. Arendt, M. S. Kaiser. Rainfall correlates of bird population fluctuations in a Puerto Rican dry forest: a nine year study. Wilson Bull., 96, 575–593 (1984).

99. P. Macario et al. Apparent survival and cost of reproduction for white-lined tanager (*Tachyphonus rufus*, Thraupidae) in the northern Atlantic Rainforest, Brazil. PloS One, 12, e0185890 (2017).

100. T. B. Ryder, T. S. Sillett. Climate, demography and lek stability in an Amazonian bird. Proc. Roy. Soc. B, 283, 20152314 (2016).

101. A. E. Jahn et al. Seasonal differences in rainfall, food availability, and the foraging behavior of tropical kingbirds in the southern Amazon basin. J. Field Ornithol., 81, 340–348 (2010).

102. R. Greenberg, J. Gradwohl. Constant density and stable territoriality in some tropical insectivorous birds. Oecologia, 69, 618–625 (1986).

103. C. H. Şekercioğlu et al. Disappearance of insectivorous birds from tropical forest fragments. Proc. Natl. Acad. Sci. 99, 263–267 (2002).

104. T. W. Sherry, C. M. Kent, N. V. Sánchez, C. H. Şekercioğlu. Insectivorous birds in the Neotropics: ecological radiations, specialization, and coexistence in species-rich communities. Auk, 137, ukaa049 (2020).

105. S. Woltmann, T. W. Sherry. High apparent annual survival and stable territory dynamics of chestnut-backed antbird (*Myrmeciza exsul*) in a large Costa Rican rain forest preserve. Wilson J. Ornithol., 123, 15–23 (2011).

106. S. R. Montanari, J. P. A. Hobbs, M. S. Pratchett, L. K. Bay, L. Van Herwerden. Does genetic distance between parental species influence outcomes of hybridization among coral reef butterflyfishes? Mol. Ecol., 23, 2757–2770 (2014).

107. N. H. Barton, G. M. Hewitt. Analysis of hybrid zones. Annu. Rev. Eco. Syst., 16, 113–148 (1985).

108. G. M. Hewitt. Hybrid zones-natural laboratories for evolutionary studies. Trends Ecol. Evol., 3, 158–167 (1988).

109. F. M. Chapman. The Distribution of Bird-Life in Colombia. (American Museum of Natural History, 1917).

110. G. R. Angehr, R. Dean. The Birds of Panama: A Field Guide. (Cornell University Press, 2010).

111. R. S. Ridgely, J. A Gwynne. A Guide to the Birds of Panama, with Costa Rica, Nicaragua, and Honduras. 2nd ed. (Princeton University Press. 1992).

112. D. C. Siegel, S. L. Olson. The Birds of the Republic of Panama, Part 5. Gazetteer and Bibliography. (Buteo Books, 2008).

113. A. Wetmore. The Birds of the Republic of Panama, Part 1. Tinamidae (Tinamous) to Rynchopidae (Skimmers). (Smithsonian Institution Press, 1965).

114. A. Wetmore. The Birds of the Republic of Panama, Part 2. Columbidae (Pigeons) to Picidae (Woodpeckers). (Smithsonian Institution Press, 1968).

115. A. Wetmore. The Birds of the Republic of Panama, Part 3. Passeriformes: Dendrocolaptidae (Woodcreepers) to Oxyruncidae (Sharpbills). (Smithsonian Institution Press, 1972).

116. A. Wetmore, R. F. Pasquier, S. L. Olson. The Birds of the Republic of Panama, Part 4. Passeriformes: Hirundinidae (Swallows) to Frigillidae (Finches). (Smithsonian Institution Press, 1984).

117. J. F. McLaughlin, J. L. Garzón, O. G. López Ch., M. J. Miller. A preliminary bird list from Río Luis, Veraguas province, provides further insight into an avian suture zone in Caribbean Panama. Cotinga, 42, 77–81 (2020).

118. Z. A. Cheviron, S. J. Hackett, A. P. Capparella. Complex evolutionary history of a Neotropical lowland forest bird (*Lepidothrix coronata*) and its implications for historical hypotheses of the origin of Neotropical avian diversity. Mol. Phylogenet. Evol., 36, 338–357 (2005).

119. I. J. Lovette. Molecular phylogeny and plumage signal evolution in a trans Andean and circum Amazonian avian species complex. Mol. Phylogenet. Evol., 32, 512–523 (2004).

120. B. Milá et al. A trans-Amazonian screening of mtDNA reveals deep intraspecific divergence in forest birds and suggests a vast underestimation of species diversity. PloS One, 7, e40541 (2012).

121. M. J. Miller et al. Demographic consequences of foraging ecology explain genetic diversification in Neotropical bird species. Ecol. Lett., 24, 563–571 (2021).

122. M. J. Miller et al. Phylogeography of the rufous-tailed hummingbird (*Amazilia tzacatl*). Condor, 113, 806–816 (2011).

123. H. Vázquez-Miranda, A. G. Navarro-Sigüenza, K. E. Omland. Phylogeography of the rufous-naped wren (*Campylorhynchus rufinucha*): speciation and hybridization in Mesoamerica. Auk, 126, 765–78 (2009).

124. J. H. Gillespie. Population Genetics: A Concise Guide. (JHU Press, 2004).

125. M. Kimura. Evolutionary rate at the molecular level. Nature, 217, 624–626 (1968).

126. M. Kimura. The Neutral Theory of Molecular Evolution. (Cambridge University Press, 1983).

127. M. Kimura, T. Ohta. On the rate of molecular evolution. J. Mol. Evol., 1, 1–17 (1971).

128. L. Bromham, D. Penny. The modern molecular clock. Nat. Rev. Genet., 4, 216–224 (2003).

129. T. Ohta. The nearly neutral theory of molecular evolution. Annu. Rev. Eco. Syst., 23, 263–286 (1992).

130. J. T. Weir, D. Schluter. Calibrating the avian molecular clock. Mol. Ecol., 17, 2321–2328 (2008).

131. A. Qvarnström, R. I. Bailey. Speciation through evolution of sex-linked genes. Heredity, 102, 4–15 (2009).

132. B. Charlesworth, J. A. Coyne, N. H. Barton. The relative rates of evolution of sex chromosomes and autosomes. Am. Nat., 130, 113–146 (1987).

133. S. Wright. Inbreeding and homozygosis. Proc. Natl. Acad. Sci., 19, 411–20 (1933).

134. D. M. Hooper, S. C. Griffith, T. D. Price. Sex chromosome inversions enforce reproductive isolation across an avian hybrid zone. Mol. Ecol., 28, 1246–1262 (2019).

135. W. R. Rice. The accumulation of sexually antagonistic genes as a selective agent promoting the evolution of reduced recombination between primitive sex chromosomes. Evolution, 41, 911–914 (1987).

136. V. B. Kaiser, H. Ellegren. Nonrandom distribution of genes with sex-biased expression in the chicken genome. Evolution, 60, 1945–1951 (2006).

137. R. Storchová, P. Divina. Nonrandom representation of sex-biased genes on the chicken Z chromosome. J. Mol. Evol. 63, 676–681 (2006).

138. J. P. McEntee, J. G. Burleigh, S. Singhal. Dispersal predicts hybrid zone widths across animal diversity: implications for species borders under incomplete reproductive isolation. Am. Nat., 196, 9–28 (2020).

139. D. P. L. Toews et al. Plumage genes and little else distinguish the genomes of hybridizing warblers. Curr. Biol., 26, 2313–2318 (2016).

140. L. Campagna et al. Repeated divergent selection on pigmentation genes in a rapid finch radiation. Sci. Adv., 3, e1602404 (2017).

141. S. Edmands. Does parental divergence predict reproductive compatibility? Trends Ecol. Evol., 17, 520–27 (2002).

142. S. Hogner et al. Deep sympatric mitochondrial divergence without reproductive isolation in the common redstart Phoenicurus phoenicurus. Ecol. Evol., 2, 2974–2988 (2012).

143. C. Dufresnes et al. Mass of genes rather than master genes underlie the genomic architecture of amphibian speciation. Proc. Natl. Acad. of Sci., 118, e2103963118 (2021).

144. N. H. Barton, G. M. Hewitt (1989). Adaptation, speciation and hybrid zones. Nature, 341, 497–503

145. Z. Gompert, E. G. Mandeville, C. A. Buerkle. Analysis of population genomic data from hybrid zones. Annu. Rev. Ecol. Evol. S., 48, 207–229 (2017).

146. G. M. Hewitt. Speciation, hybrid zones and phylogeography—or seeing genes in space and time. Mol. Ecol., 10, 537–549 (2001).

147. M. B. Bush, P. A. Colinvaux (1990). A pollen record of a complete glacial cycle from lowland Panama. J. Veg. Sci., 1, 105–118

148. D. R. Piperno, M. B. Bush, P. A. Colinvaux, Paleoecological perspectives on human adaptation in central Panama, II: the Holocene. Geoarchaeology, 6, 227–250 (1991).

149. J. F. McLaughlin et al. Comparative phylogeogeography reveals widespread cryptic diversity driven by ecology in Panamanian birds. [preprint?] (2022).

150. M. G. Harvey, B. T. Smith, T. C. Glenn, B. C. Faircloth, R. T. Brumfield Sequence capture versus restriction site associated DNA sequencing for shallow systematics. Syst. Biol., 65, 910–924 (2016).

151. B. T. Smith et al. The drivers of tropical speciation. Nature, 515, 406–409 (2014).

152. B. C. Faircloth McCormack, J. E., Crawford, N. G., M. G. Harvey, R. T. Brumfield, T. C. Glenn. Ultraconserved elements anchor thousands of genetic markers spanning multiple evolutionary timescales. Syst. Biol., 61, 717–726 (2012).

153. T. C. Glenn et al. Adapterama I: universal stubs and primers for 384 unique dual-indexed or 147,456 combinatorially-indexed Illumina libraries (iTru, iNext). PeerJ, 7, e7755 (2019).

154. A. M. Bolger, M. Lohse, B. Usadel. Trimmomatic: a flexible trimmer for Illumina sequence data. Bioinformatics, 30, 2114–2120 (2014).

155. B. C. Faircloth. 2013. “Illumiprocessor: A Trimmomatic Wrapper for Parallel Adapter and Quality Trimming.” See Http://dx. Doi. org/10. 6079/J9ILL (accessed 4 November 2016).

156. Bushnell, Brian. 2014. “BBMap: A Fast, Accurate, Splice-Aware Aligner.” LBNL-7065E. Lawrence Berkeley National Lab. (LBNL), Berkeley, CA (United States).

157. M. G. Grabherr et al. Full-length transcriptome assembly from RNA-seq data without a reference genome. Nat. Biotechnol., 29, 644–652 (2011).

158. B. C. Faircloth. PHYLUCE is a software package for the analysis of conserved genomic loci. Bioinformatics, 32, 786–788 (2016).

159. A. McKenna et al. The Genome Analysis Toolkit: a MapReduce framework for analyzing next-generation DNA sequencing data. Genome Res., 20, 1297–1303 (2010).

160. H. Li. 2013. “Aligning Sequence Reads, Clone Sequences and Assembly Contigs with BWA-MEM.” arXiv [q-bio. GN]. arXiv. http://arxiv.org/abs/1303.3997.

161. H. Li, R. Durbin. Fast and accurate short read alignment with Burrows–Wheeler transform. Bioinformatics, 25, 1754–1760 (2009).

162. H. Li et al. The sequence alignment/map format and SAMtools. Bioinformatics, 25, 2078–2079 (2009).

163. Broad Institute. (2019). “Picard Toolkit.” http://broadinstitute.github.io/picard/.

164. P. Danecek et al. The variant call format and VCFtools. Bioinformatics, 27, 2156–2158 (2011).

165. Z. Zhang, S. Schwartz, L. Wagner, W. Miller. A greedy algorithm for aligning DNA sequences. J.Comput. Biol., 7, 203–214 (2000).

166. N. Dierckxsens, P. Mardulyn, G. Smits. NOVOPlasty: de novo assembly of organelle genomes from whole genome data. Nucleic Acids Res., 45, e18 (2017).

167. T. Jombart, I. Ahmed. Adegenet 1.3-1: new tools for the analysis of genome-wide SNP data. Bioinformatics, 27, 3070–3071 (2011).

168. J. K. Pritchard, M., Stephens, P. Donnelly. Inference of population structure using multilocus genotype data. Genetics, 155, 945–959 (2000).

169. M. Jakobsson, N. A. Rosenberg. CLUMPP: a cluster matching and permutation program for dealing with label switching and multimodality in analysis of population structure. Bioinformatics, 23, 1801–1806 (2007).

170. D. A. Earl, B. M. vonHoldt. STRUCTURE HARVESTER: a website and program for visualizing STRUCTURE output and implementing the Evanno method. Conserv. Genet. Resour., 4, 359–361 (2012).

171. G. Evanno, S. Regnaut, J. Goudet. Detecting the number of clusters of individuals using the software STRUCTURE: a simulation study. Mol. Ecol., 14, 2611–2620 (2005).

172. N. A. Rosenberg et al. Genetic structure of human populations. Science, 298, 2381–2385 (2002).

173. J. R. Karr. Seasonality, resource availability, and community diversity in tropical bird communities. Am. Nat., 110, 973–994 (1976).

174. D. J. Levey, F. G. Stiles. “Birds: ecology, behavior, and taxonomic affinities” in La Selva: Ecology and Natural History of a Neotropical Rain Forest, L. A. McDade, K. S. Bawa, H. A. Hespenheide, G. A. Hartshorn, Eds. (University of Chicago Press, 1994), pp 217–228.

