## Supplemental Tables and FIgures for "Time in allopatry does not predict the outcome of secondary contact in lowland Panamanian birds"

**Table S1:** Sequencing and UCE recovery results. UCE= UCE enriched sample, WGS= unenriched whole genome shotgun library. Museum numbers provided where specimens have been fully cataloged, with field catalog numbers provided for samples which have not been added to museum databases. CUMV: Cornell University Museum of Vertebrates, FMNH: Field Museum of Natural History, STRIBC: Smithsonian Tropical Research Institute Bird Collection, UAM: University of Alaska Museum, UWBM: University of Washington Burke Museum.

| **Taxon** | **Museum Number** | **Catalog Number** | **Population** | **Type** | **Reads (millions)** | **SRA Accession Number** |
| --- | --- | --- | --- | --- | --- | --- |
| ***Arremon aurantiirostris*** | UWBM93635 |  | Honduras: Copán | WGS | 82.8 | SAMN28920643 |
| *Arremon aurantiirostris* | UWBM56060 |  | Nicaragua: La Luz | WGS | 120.6 | SAMN28920644 |
| *Arremon aurantiirostris* | UWBM70031 |  | Nicaragua: La Luz | WGS | 104.4 | SAMN28920645 |
| *Arremon aurantiirostris* |  | MJM2298 | Panama: Bocas del Toro, Bosque Protector de Palo Seco | WGS | 47.4 | SAMN28920646 |
| *Arremon aurantiirostris* | STRIBC5367 | MJM2299 | Panama: Bocas del Toro, Bosque Protector de Palo Seco | WGS | 42.8 | SAMN28920647 |
| *Arremon aurantiirostris* | STRIBC2287 | MJM2930 | Panama: Bocas del Toro, Changuinola | UCE |  | SAMN28920648 |
| *Arremon aurantiirostris* | STRIBC2266 | MJM4225 | Panama: Bocas del Toro, Changuinola | UCE |  | SAMN28920649 |
| *Arremon aurantiirostris* |  | JTW288 | Panama: Comarca Ngäbe-Buglé, Cerro Chalite | UCE |  | SAMN28920650 |
| *Arremon aurantiirostris* | FMNH470780 | GMS2011 | Panama: Bocas del Toro, Bosque Protector de Palo Seco | UCE |  | SAMN28920651 |
| *Arremon aurantiirostris* |  | JFM022 | Panama: Veraguas, Río Luis | WGS | 72.8 | SAMN28920652 |
| *Arremon aurantiirostris* |  | JFM037 | Panama: Veraguas, Río Luis | WGS | 203.8 | SAMN28920653 |
| *Arremon aurantiirostris* |  | JFM044 | Panama: Veraguas, Río Luis | WGS | 40.8 | SAMN28920654 |
| *Arremon aurantiirostris* |  | JFM047 | Panama: Veraguas, Río Luis | WGS | 126.4 | SAMN28920655 |
| *Arremon aurantiirostris* | UWBM108325 | JMD114 | Panama: Veraguas, Santa Fé | UCE | 4.68 | SAMN28920656 |
| *Arremon aurantiirostris* | STRIBC2762 | MJM6862 | Panama: Veraguas, Santa Fé | UCE | 0.42 | SAMN28920657 |
| *Arremon aurantiirostris* | STRIBC2577 | MJM6871 | Panama: Veraguas, Santa Fé | UCE | 1.12 | SAMN28920658 |
| *Arremon aurantiirostris* | STRIBC2811 | MJM6932 | Panama: Veraguas, Santa Fé | UCE | 3.35 | SAMN28920659 |
| *Arremon aurantiirostris* | STRIBC4312 | MJM8129 | Panama: Coclé, La Pintada, Coclesito | UCE | 0.85 | SAMN28920660 |
| *Arremon aurantiirostris* | STRIBC3568 | MJM8186 | Panama: Coclé, La Pintada, Coclesito | UCE | 4.87 | SAMN28920661 |
| *Arremon aurantiirostris* | UWBM106601 | GMS1179 | Panama: Colón, Achiote | UCE | 0.60 | SAMN28920662 |
| *Arremon aurantiirostris* | UWBM106602 | GMS1180 | Panama: Colón, Achiote | UCE | 0.67 | SAMN28920663 |
| *Arremon aurantiirostris* | STRIBC2810 | MJM6628 | Panama: Coclé, San Juan, Aguas Claras | UCE | 0.73 | SAMN28920664 |
| *Arremon aurantiirostris* | STRIBC2268 | MJM7004 | Panama: Coclé, San Juan, Aguas Claras | UCE | 3.33 | SAMN28920665 |
| *Arremon aurantiirostris* | UWBM76885 | RCF2020 | Panama: Panamá, Panama City, confluence of Rios Chagres and Chagrecito | UCE | 0.66 | SAMN28920666 |
| *Arremon aurantiirostris* | UWBM76954 | SMB205 | Panama: Panamá, Panama City, confluence of Rios Chagres and Chagrecito | UCE | 3.96 | SAMN28920667 |
| *Arremon aurantiirostris* | STRIBC3944 | MJM8562 | Panama: Panama, Chiman, Cerro Chucanti | UCE | 0.96 | SAMN28920668 |
| *Arremon aurantiirostris* | STRIBC3713 | MJM8563 | Panama: Panama, Chiman, Cerro Chucanti | UCE | 3.20 | SAMN28920669 |
| *Arremon aurantiirostris* | UAM22809 | MJM1933 | Panama: Darién, Cana | WGS | 64.3 | SAMN28920670 |
| *Arremon aurantiirostris* | UAM25715 | MJM1949 | Panama: Darién, Cana | UCE | 0.65 | SAMN28920671 |
| *Arremon aurantiirostris* | UAM22803 | MJM1979 | Panama: Darién, Cana | UCE | 0.75 | SAMN28920672 |
| *Arremon aurantiirostris* | UAM22807 | MJM2005 | Panama: Darién, Cana | UCE | 0.61 | SAMN28920673 |
| *Arremon aurantiirostris* | UAM25716 | MJM1931 | Panama: Darién, Cana | UCE | 0.68 | SAMN28920674 |
| ***Cantorchilus nigricapillus*** | STRIBC1447 | MJM2656 | Panama: Bocas del Toro, Isla Escudo de Veraguas | WGS | 229.4 | SAMN28920675 |
| *Cantorchilus nigricapillus* |  | JFM052 | Panama: Veraguas, Río Luis | WGS | 226.8 | SAMN28920676 |
| *Cantorchilus nigricapillus* |  | JFM053 | Panama: Veraguas, Río Luis | WGS | 40.4 | SAMN28920677 |
| *Cantorchilus nigricapillus* | STRIBC3992 |  | Panama: Bocas del Toro, Isla Escudo de Veraguas | WGS | 123.6 | SAMN28920678 |
| *Cantorchilus nigricapillus* | STRIBC3982 |  | Panama: Bocas del Toro, Isla Escudo de Veraguas | WGS | 34.3 | SAMN28920679 |
| *Cantorchilus nigricapillus* | STRIBC1474 | MJM2759 | Panama: Colón, Achiote | UCE | 3.8 | SAMN28920680 |
| *Cantorchilus nigricapillus* | STRIBC1476 | MJM4664 | Panama: Colón, Achiote | UCE | 4.0 | SAMN28920681 |
| *Cantorchilus nigricapillus* |  | PA-TNI472 | Panama: Panamá, Cerro Azul | UCE | 0.50 | SAMN28920682 |
| *Cantorchilus nigricapillus* | STRIBC4216 |  | Panama: Colón, Santa Isabel, Palenque | WGS | 127.4 | SAMN28920683 |
| *Cantorchilus nigricapillus* |  | PA-TNI26392 | Panama: Panamá, Serranía de San Blas, west end | UCE | 0.71 | SAMN28920684 |
| *Cantorchilus nigricapillus* |  | PA-TNI26393 | Panama: Panamá, Serranía de San Blas, west end | UCE | 0.68 | SAMN28920685 |
| *Cantorchilus nigricapillus* |  | PA-TNI1906 | Panama: Panama, Chiman, Cerro Chucanti | UCE | 0.66 | SAMN28920686 |
| *Cantorchilus nigricapillus* | PA-TNI666 |  | Panama: Darién, Puerto Piña | WGS | 211.7 | SAMN28920687 |
| *Cantorchilus nigricapillus* | PA-TNI46569 |  | Panama: Darién, Rancho Frío | WGS | 78.7 | SAMN28920688 |
| *Cantorchilus nigricapillus* | STRIBC4840 |  | Panama: Darién, Rancho Frío | WGS | 51.9 | SAMN28920689 |
| *Cantorchilus nigricapillus* | STRIBC4841 |  | Panama: Darién, Rancho Frío | WGS | 96.9 | SAMN28920690 |
| *Cantorchilus nigricapillus* | UAM25132 | JMM1041 | Panama: Darién, Cana | UCE | 1.00 | SAMN28920691 |
| *Cantorchilus nigricapillus* | UAM31187 | JMM1042 | Panama: Darién, Cana | WGS | 60.1 | SAMN28920692 |
| *Cantorchilus nigricapillus* | UAM31480 | JMM1043 | Panama: Darién, Cana | WGS | 82.9 | SAMN28920693 |
| *Cantorchilus nigricapillus* | UAM27613 | KSW4884 | Panama: Darién, Cana | UCE | 0.90 | SAMN28920694 |
| *Cantorchilus nigricapillus* |  | MJM2125 | Panama: Darién, Cana | UCE | 0.53 | SAMN28920695 |
| *Cantorchilus nigricapillus* |  | EC-TNI2047 | Ecuador: Pichincha, Mindo | WGS | 189.4 | SAMN28920696 |
| *Cantorchilus nigricapillus* |  | EC-TNI12053 | Ecuador: Pichincha, Mindo | WGS | 111.0 | SAMN28920697 |
| ***Cyanocompsa cyanoides*** | UAM24530 | ABJ481 | Belize: Toledo, Big Falls | UCE | 3.7 | SAMN28920698 |
| *Cyanocompsa cyanoides* | UAM18351 | ABJ865 | Belize: Toledo, Big Falls | UCE | 0.7 | SAMN28920699 |
| *Cyanocompsa cyanoides* | UAM15286 | KSW3847 | Belize: Toledo, Big Falls | UCE | 5.0 | SAMN28920700 |
| *Cyanocompsa cyanoides* | UWBM70035 | DAB1176 | Nicaragua: La Luz | WGS | 100.6 | SAMN28920701 |
| *Cyanocompsa cyanoides* | UWBM56337 | DAB1256 | Nicaragua: La Luz | WGS | 93.6 | SAMN28920702 |
| *Cyanocompsa cyanoides* | STRIBC2421 | MJM3014 | Panama: Bocas del Toro, Changuinola | UCE | 5.5 | SAMN28920703 |
| *Cyanocompsa cyanoides* | STRIBC2420 | MJM4149 | Panama: Bocas del Toro, Changuinola | UCE | 5.6 | SAMN28920704 |
| *Cyanocompsa cyanoides* | STRIBC2422 | MJM3016 | Panama: Bocas del Toro, Changuinola | UCE | 5.0 | SAMN28920705 |
| *Cyanocompsa cyanoides* | CUMV51040 | IJL04191 | Panama: Bocas del Toro, Chiriquí Grande, Rio La Gloria | UCE | 4.5 | SAMN28920706 |
| *Cyanocompsa cyanoides* | STRIBC2435 | MJM2337 | Panama: Comarca Ngäbe-Buglé, Cerro Chalite | UCE | 6.0 | SAMN28920707 |
| *Cyanocompsa cyanoides* | STRIBC2423 | MJM6963 | Panama: Veraguas, Santa Fé | UCE | 5.0 | SAMN28920708 |
| *Cyanocompsa cyanoides* | UWBM111233 | JK04142 | Panama: Veraguas, Santa Fé | UCE | 5.4 | SAMN28920709 |
| *Cyanocompsa cyanoides* | STRIBC2428 | MJM6699 | Panama: Colón, Achiote | UCE | 4.1 | SAMN28920710 |
| *Cyanocompsa cyanoides* | STRIBC2438 | MJM6683 | Panama: Colón, Achiote | UCE | 6.1 | SAMN28920711 |
| *Cyanocompsa cyanoides* | STRIBC3390 | MJM7996 | Panama: Colón, Achiote | UCE | 5.1 | SAMN28920712 |
| ***Henicorhina leucosticta*** | UWBM123523 | JK06130 | Panama: Bocas del Toro, Changuinola | UCE | 0.37 | SAMN28920713 |
| *Henicorhina leucosticta* | UWBM123380 | JMD754 | Panama: Bocas del Toro, Changuinola | UCE | 0.44 | SAMN28920714 |
| *Henicorhina leucosticta* | UWBM123642 | JK06125 | Panama: Bocas del Toro, Changuinola | UCE | 0.58 | SAMN28920715 |
| *Henicorhina leucosticta* |  | JTW280 | Panama: Comarca Ngäbe-Buglé, Cerro Chalite | UCE | 1.34 | SAMN28920716 |
| *Henicorhina leucosticta* | UWBM106448 | GMS1021 | Panama: Veraguas, Santa Fé | UCE | 0.88 | SAMN28920717 |
| *Henicorhina leucosticta* | STRIBC1528 | MJM6908 | Panama: Veraguas, Santa Fé | UCE | 0.54 | SAMN28920718 |
| *Henicorhina leucosticta* | UWBM111225 | JK04134 | Panama: Veraguas, Santa Fé | UCE | 1.12 | SAMN28920719 |
| *Henicorhina leucosticta* | STRIBC1534 | MJM3373 | Panama: Coclé, La Pintada, El Copé | UCE | 0.35 | SAMN28920720 |
| *Henicorhina leucosticta* | STRIBC1531 | MJM3463 | Panama: Coclé, La Pintada, El Copé | UCE | 0.57 | SAMN28920721 |
| *Henicorhina leucosticta* | UAM24660 | MJM1420 | Panama: Colón, Achiote | UCE | 1.11 | SAMN28920722 |
| *Henicorhina leucosticta* | STRIBC1536 | MJM4504 | Panama: Colón, Achiote | UCE | 0.94 | SAMN28920723 |
| *Henicorhina leucosticta* | UAM24661 | MJM1044 | Panama: Panamá, Cerro Azul | UCE | 0.32 | SAMN28920724 |
| *Henicorhina leucosticta* | UAM24580 | JMM907 | Panama: Panamá, Cerro Azul | UCE | 0.61 | SAMN28920725 |
| *Henicorhina leucosticta* | UAM22726 | MJM696 | Panama: Panamá, Cerro Jefe | UCE | 0.23 | SAMN28920726 |
| *Henicorhina leucosticta* | UAM22728 | MJM1057 | Panama: Panamá, Cerro Jefe | UCE | 0.92 | SAMN28920727 |
| *Henicorhina leucosticta* | UWBM107228 | GMS1830 | Panama: Panamá, Lago Bayano | UCE | 0.22 | SAMN28920728 |
| *Henicorhina leucosticta* | UWBM120908 | JMD657 | Panama: Panama, Chiman, Cerro Chucanti | UCE | 1.04 | SAMN28920729 |
| *Henicorhina leucosticta* | UWBM120904 | GMS1913 | Panama: Panama, Chiman, Cerro Chucanti | UCE | 0.83 | SAMN28920730 |
| *Henicorhina leucosticta* | FMNH470759 | JMD664 | Panama: Panama, Chiman, Cerro Chucanti | UCE | 0.75 | SAMN28920731 |
| *Henicorhina leucosticta* | UAM22767 | MJM2114 | Panama: Darién, Cana | UCE | 0.62 | SAMN28920732 |
| *Henicorhina leucosticta* | UAM22766 | MJM2113 | Panama: Darién, Cana | UCE | 0.53 | SAMN28920733 |
| *Henicorhina leucosticta* | UAM24008 | MJM2089 | Panama: Darién, Cana | UCE | 0.22 | SAMN28920734 |
| *Henicorhina leucosticta* | UAM22761 | MJM1987 | Panama: Darién, Cana | UCE | 0.16 | SAMN28920735 |
| ***Malacoptila panamensis*** | STRIBC2788 | MJM4404 | Panama: Bocas del Toro, Changuinola | UCE | 5.0 | SAMN28920736 |
| *Malacoptila panamensis* | STRIBC2786 | MJM4298 | Panama: Bocas del Toro, Changuinola | UCE | 4.4 | SAMN28920737 |
| *Malacoptila panamensis* | STRIBC0500 | MJM3097 | Panama: Bocas del Toro, Changuinola | UCE | 5.2 | SAMN28920738 |
| *Malacoptila panamensis* | STRIBC2787 | MJM4308 | Panama: Bocas del Toro, Changuinola | UCE | 4.0 | SAMN28920739 |
| *Malacoptila panamensis* |  | JTW318 | Panama: Comarca Ngäbe-Buglé, Cerro Chalite | UCE | 2.5 | SAMN28920740 |
| *Malacoptila panamensis* | STRIBC7888 | JFM074 | Panama: Veraguas, Río Luis | WGS | 81.6 | SAMN28920741 |
| *Malacoptila panamensis* | UAM20359 | MJM265 | Panama: Coclé, La Pintada, Coclesito | UCE | 4.3 | SAMN28920742 |
| *Malacoptila panamensis* | STRIBC3597 | MJM8074 | Panama: Coclé, La Pintada, Coclesito | UCE | 5.0 | SAMN28920743 |
| *Malacoptila panamensis* | UAM20417 | MJM324 | Panama: Coclé, La Pintada, Coclesito | UCE | 5.3 | SAMN28920744 |
| *Malacoptila panamensis* | STRIBC2564 | MJM2819 | Panama: Colón, Achiote | UCE | 6.4 | SAMN28920745 |
| *Malacoptila panamensis* | STRIBC2563 | MJM2822 | Panama: Colón, Achiote | UCE | 4.3 | SAMN28920746 |
| *Malacoptila panamensis* | STRIBC0503 | MJM4485 | Panama: Colón, Achiote | UCE | 5.1 | SAMN28920747 |
| *Malacoptila panamensis* | STRIBC2908 | MJM7208 | Panama: Colón, Gamboa | UCE | 4.3 | SAMN28920748 |
| *Malacoptila panamensis* | FMNH470655 | JMD697 | Panama: Panama, Chiman, Cerro Chucanti | UCE | 3.7 | SAMN28920749 |
| *Malacoptila panamensis* | FMNH470657 | JMD695 | Panama: Panama, Chiman, Cerro Chucanti | UCE | 4.2 | SAMN28920750 |
| *Malacoptila panamensis* | FMNH470656 | JMD696 | Panama: Panama, Chiman, Cerro Chucanti | UCE | 4.8 | SAMN28920751 |
| *Malacoptila panamensis* | STRIBC4814 | MJM9375 | Panama: Darién, Rancho Frío | UCE | 4.1 | SAMN28920752 |
| *Malacoptila panamensis* |  | B17539 | Panama: Darién, Rancho Frío | UCE | 4.4 | SAMN28920753 |
| *Malacoptila panamensis* | STRIBC4836 | MJM9397 | Panama: Darién, Rancho Frío | UCE | 4.3 | SAMN28920754 |
| ***Myrmeciza exsul*** |  | B58101 | Almirante | WGS | 79.3 | SAMN28920755 |
| *Myrmeciza exsul* | STRIBC 0825 | MJM4263 | Panama: Bocas del Toro, Changuinola | WGS | 105.6 | SAMN28920756 |
| *Myrmeciza exsul* | STRIBC4141 |  | Panama: Coclé, La Pintada, Coclesito | WGS | 118.3 | SAMN28920757 |
| *Myrmeciza exsul* | STRIBC0796 |  | Panama: Colón, Achiote | WGS | 55.3 | SAMN28920758 |
| *Myrmeciza exsul* | STRIBC 2961 | MJM7205 | Panama: Colón, Gamboa | WGS | 59.2 | SAMN28920759 |
| *Myrmeciza exsul* | STRIBC 3856 | MJM8033 | Panama: Colón, Gamboa | WGS | 49.9 | SAMN28920760 |
| *Myrmeciza exsul* |  | GKD256 | Panama: Panamá, Panama City, confluence of Rios Chagres and Chagrecito | WGS | 99.4 | SAMN28920761 |
| *Myrmeciza exsul* |  | RCF2031 | Panama: Panamá, Panama City, confluence of Rios Chagres and Chagrecito | WGS | 145.5 | SAMN28920762 |
| *Myrmeciza exsul* |  | SMB223 | Panama: Panamá, Panama City, confluence of Rios Chagres and Chagrecito | WGS | 61.0 | SAMN28920763 |
| *Myrmeciza exsul* |  | MJM0503 | Panama: Panamá, Cerro Azul | WGS | 88.2 | SAMN28920764 |
| *Myrmeciza exsul* | STRIBC 0799 | MJM5636 | Panama: Panamá, Cerro Azul | WGS | 90.1 | SAMN28920765 |
| *Myrmeciza exsul* | STRIBC 3921 | MJM8564 | Panama: Panama, Chiman, Cerro Chucanti | WGS | 74.6 | SAMN28920766 |
| *Myrmeciza exsul* | STRIBC 3742 | MJM8565 | Panama: Panama, Chiman, Cerro Chucanti | WGS | 89.9 | SAMN28920767 |
| *Myrmeciza exsul* | STRIBC3538 |  | Panama: Darién, Chepigana, Chucanaque, El Salto | WGS | 140.5 | SAMN28920768 |
| *Myrmeciza exsul* | STRIBC 4140 | MJM9020 | Panama: Comarca Emberá-Wounann, Cémaco, Peña Bijagual | WGS | 96.8 | SAMN28920769 |
| *Myrmeciza exsul* | STRIBC 4389 | MJM9148 | Panama: Comarca Emberá-Wounann, Cémaco, Peña Bijagual | WGS | 153.8 | SAMN28920770 |
| *Myrmeciza exsul* |  | MJM985 | Tropic Star | WGS | 77.2 | SAMN28920771 |
| *Myrmeciza exsul* | STRIBC4835 |  | Panama: Darién, Rancho Frío | WGS | 111.8 | SAMN28920772 |
| *Myrmeciza exsul* | STRIBC4837 |  | Panama: Darién, Rancho Frío | WGS | 102.7 | SAMN28920773 |
| *Myrmeciza exsul* | MJM2023 |  | Panama: Darién, Cana | WGS | 46.5 | SAMN28920774 |
| ***Pachysylvia decurtata*** | STRIBC1422 | MJM6350 | Panama: Bocas del Toro, Bosque Protector de Palo Seco | UCE | 4.5 | SAMN28920775 |
| *Pachysylvia decurtata* | STRIBC1423 | B17498 | Panama: Bocas del Toro, Bosque Protector de Palo Seco | UCE | 4.4 | SAMN28920776 |
| *Pachysylvia decurtata* | UWBM111263 | JK04172 | Panama: Veraguas, Santa Fé | UCE | 4.1 | SAMN28920777 |
| *Pachysylvia decurtata* | UWBM111255 | JK04164 | Panama: Veraguas, Santa Fé | UCE | 5.2 | SAMN28920778 |
| *Pachysylvia decurtata* | UWBM106463 | GMS1036 | Panama: Veraguas, Santa Fé | UCE | 4.1 | SAMN28920779 |
| *Pachysylvia decurtata* | STRIBC1423 | MJM2660 | Panama: Panamá, Cerro Azul | UCE | 5.1 | SAMN28920780 |
| *Pachysylvia decurtata* | UWBM111193 | JK04100 | Panama: Panamá, Cerro Jefe | UCE | 4.0 | SAMN28920781 |
| *Pachysylvia decurtata* |  | JMM910 | Panama: Panamá, Cerro Azul | UCE | 4.7 | SAMN28920782 |
| *Pachysylvia decurtata* | UWBM108154 | GMS983 | Panama: Panamá, Cerro Jefe | UCE | 4.0 | SAMN28920783 |
| *Pachysylvia decurtata* | STRIBC3764 | MJM8573 | Panama: Panama, Chiman, Cerro Chucanti | UCE | 3.5 | SAMN28920784 |
| *Pachysylvia decurtata* | UWBM112234 | JK06044 | Panama: Panama, Chiman, Cerro Chucanti | UCE | 3.5 | SAMN28920785 |
| *Pachysylvia decurtata* | UWBM108897 | JMD719 | Panama: Panama, Chiman, Cerro Chucanti | UCE | 4.6 | SAMN28920786 |
| *Pachysylvia decurtata* |  | B46582 | Panama: Darién, Rancho Frío | UCE | 4.2 | SAMN28920787 |
| *Pachysylvia decurtata* |  | B46600 | Panama: Darién, Rancho Frío |  |  | SAMN28920788 |
| ***Ramphocelus passerinii*** |  | PA-RPA46461 | Panama: Bocas del Toro, Guabito | WGS | 90.0 | SAMN28920789 |
| *Ramphocelus passerinii* | STRIBC4562 | MJM4054 | Panama: Bocas del Toro, Changuinola | WGS | 70.8 | SAMN28920790 |
| *Ramphocelus passerinii* | STRIBC1898 | MJM4007 | Panama: Bocas del Toro, Changuinola | WGS | 71.7 | SAMN28920791 |
| *Ramphocelus passerinii* | STRIBC1901 | MJM4146 | Panama: Bocas del Toro, Changuinola | WGS | 97.0 | SAMN28920792 |
| *Ramphocelus passerinii* | STRIBC4629 | MJM4170 | Panama: Bocas del Toro, Changuinola | WGS | 63.9 | SAMN28920793 |
| *Ramphocelus passerinii* |  | PA-RPA46444 | Panama: Bocas del Toro, Chiriquí Grande | WGS | 98.5 | SAMN28920794 |
| *Ramphocelus passerinii* |  | PA-RPA46475 | Panama: Bocas del Toro, Chiriquí Grande | WGS | 70.1 | SAMN28920795 |
| ***Ramphocelus flammigerus*** | STRIBC3623 | MJM8197 | Panama: Coclé, La Pintada, Coclesito | WGS | 53.6 | SAMN28920796 |
| *Ramphocelus flammigerus* | STRIBC7345 | MJM8132 | Panama: Coclé, La Pintada, Coclesito | WGS | 134.0 | SAMN28920797 |
| *Ramphocelus flammigerus* | STRIBC1915 | MJM2437 | Panama: Colón, Achiote | WGS | 89.6 | SAMN28920798 |
| *Ramphocelus flammigerus* | UWBM106617 | GMS1195 | Panama: Colón, Achiote | WGS | 100.3 | SAMN28920799 |
| *Ramphocelus flammigerus* | STRIBC1918 | MJM2406 | Panama: Colón, Achiote | WGS | 76.6 | SAMN28920800 |
| *Ramphocelus flammigerus* | STRIBC4280 | MJM8966 | Panama: Colón, Santa Isabel, Palenque | WGS | 70.2 | SAMN28920801 |
| *Ramphocelus flammigerus* | STRIBC7584 | MJM8967 | Panama: Colón, Santa Isabel, Palenque | WGS | 77.1 | SAMN28920802 |
| *Ramphocelus flammigerus* | STRIBC4278 | MJM9030 | Panama: Comarca Emberá-Wounann, Cémaco, Peña Bijagual | WGS | 75.4 | SAMN28920803 |
| *Ramphocelus flammigerus* | UAM34480 | KSW4857 | Panama: Darién, Cana | WGS |  | SAMN28920804 |
| *Ramphocelus flammigerus* | UAM31175 | JMM1079 | Panama: Darién, Cana | WGS | 90.5 | SAMN28920805 |
| *Ramphocelus flammigerus* | UAM25724 | JMM1102 | Panama: Darién, Cana | WGS | 118.5 | SAMN28920806 |
| ***Schiffornis veraepacis*** |  | RCF3 | Panama: Bocas del Toro, Changuinola |  | 105.6 | SAMN28920807 |
| *Schiffornis veraepacis* | STRIBC1223 | MJM3356 | Panama: Coclé, La Pintada, El Copé | WGS | 45.5 | SAMN28920808 |
| *Schiffornis veraepacis* | STRIBC1224 | MJM3495 | Panama: Coclé, La Pintada, El Copé | WGS | 78.1 | SAMN28920809 |
| *Schiffornis veraepacis* | UWBM106535 | GMS1112 | Panama: Coclé, El Valle | WGS | 70.8 | SAMN28920810 |
| *Schiffornis veraepacis* | UWBM76978 | SMB229 | Panama: Panamá, Panama City, confluence of Rios Chagres and Chagrecito | WGS | 72.5 | SAMN28920811 |
| *Schiffornis veraepacis* | UWBM108416 | JMD208 | Panama: Panamá, Cerro Azul | WGS | 99.4 | SAMN28920812 |
| *Schiffornis veraepacis* | UWBM108283 | JMD085 | Panama: Panamá, Cerro Azul | WGS | 78.7 | SAMN28920813 |
| ***Schiffornis stenorhyncha*** | STRIBC1228 | MJM5753 | Panama: Panamá, Lago Bayano | WGS | 126.8 | SAMN28920814 |
| *Schiffornis stenorhyncha* | STRIBC4338 |  | Panama: Comarca Emberá-Wounann, Cémaco, Peña Bijagual | WGS | 112.8 | SAMN28920815 |
| *Schiffornis stenorhyncha* | STRIBC3075 | MJM7345 | Panama: Darién, Chepigana, Aruza Abajo | WGS | 76.2 | SAMN28920816 |
| *Schiffornis stenorhyncha* | STRIBC3600 | MJM7878 | Panama: Darién, Chepigana, Aruza Abajo | WGS | 105.1 | SAMN28920817 |
| ***Xenops minutus*** | STRIBC0576 | MJM2904 | Panama: Bocas del Toro, Changuinola | UCE | 4.5 | SAMN28920818 |
| *Xenops minutus* | STRIBC0577 | MJM3110 | Panama: Bocas del Toro, Changuinola | UCE | 4.5 | SAMN28920819 |
| *Xenops minutus* | STRIBC0575 | MJM4121 | Panama: Bocas del Toro, Changuinola | UCE | 7.4 | SAMN28920820 |
| *Xenops minutus* | STRIB0578 | MJM4374 | Panama: Bocas del Toro, Changuinola | UCE | 4.6 | SAMN28920821 |
| *Xenops minutus* | STRIBC7889 | JFM012 | Panama: Veraguas, Río Luis | WGS | 77.8 | SAMN28920822 |
| *Xenops minutus* | STRIBC0579 | MJM3443 | Panama: Coclé, La Pintada, El Copé | UCE | 3.9 | SAMN28920823 |
| *Xenops minutus* | STRIBC0580 | MJM3525 | Panama: Coclé, La Pintada, El Copé | UCE | 3.6 | SAMN28920824 |
| *Xenops minutus* | STRIBC0586 | MJM4675 | Panama: Colón, Achiote | UCE | 3.8 | SAMN28920825 |
| *Xenops minutus* | STRIBC0583 | MJM5323 | Panama: Colón, Achiote | UCE | 4.9 | SAMN28920826 |
| *Xenops minutus* | STRIBC0584 | MJM5612 | Panama: Panamá, Cerro Azul | UCE | 4.5 | SAMN28920827 |
| *Xenops minutus* | STRIBC0589 | MJM5646 | Panama: Panamá, Cerro Azul | UCE | 4.5 | SAMN28920828 |
| *Xenops minutus* | STRIBC0581 | MJM5735 | Panama: Panamá, Lago Bayano | UCE | 5.4 | SAMN28920829 |
| *Xenops minutus* | UWBM107189 | GMS1758 | Panama: Panamá, Lago Bayano | UCE | 5.3 | SAMN28920830 |
| *Xenops minutus* | UAM36707 | JMM1012 | Panama: Darién, Cana | UCE | 3.9 | SAMN28920831 |
| *Xenops minutus* | UAM36654 | KSW4789 | Panama: Darién, Cana | UCE | 4.1 | SAMN28920832 |
| *Xenops minutus* | UAM36653 | MJM2045 | Panama: Darién, Cana | UCE | 4.2 | SAMN28920643 |
| *Xenops minutus* | UAM24576 | MJM2051 | Panama: Darién, Cana | UCE | 4.2 | SAMN28920644 |

**Table S2:** Distances used for clinal analyses. Localities blank if taxon not sampled at that site. For each, the start of the transect is indicated with a bold “0”, with subsequent distances measured from there. All distances in straight-line kilometers.

| **Location** | **Coordinates** | ***Arremon*** | ***Cantorcilus*** | ***Cyanocompsa*** | ***Henicorhina*** | ***Malacoptila*** | ***Myrmeciza*** | ***Pachysylvia*** | ***Ramphocelus*** | ***Sciffornis*** | ***Xenops*** |
| --- | --- | --- | --- | --- | --- | --- | --- | --- | --- | --- | --- |
| La Luz, Nicaragua | 13.702, -84.854 | **0** |  | **0** |  |  |  |  |  |  |  |
| Guabito | 9.4733, -82.565 |  |  |  |  |  |  |  | **0** |  |  |
| Rio Changuinola | 9.133, -82.501 | 258.59 | **0** | 258.59 | **0** | **0** | **0** |  | 7.04 | **0** | **0** |
| Almirante | 9.307,-82.423 |  |  |  |  |  | 8.65 |  |  |  |  |
| Rio La Gloria | 8.9844, -82.233 |  |  | 288.09 |  |  |  |  |  |  |  |
| Palo Seco | 8.7934, -82.189 | 292.925 |  |  |  |  |  | **0** |  |  |  |
| Chiriqui Grande | 8.7934, -82.18885 |  |  |  |  |  |  |  | 41.37 |  |  |
| Cerro Chalite | 8.858611, 82.06055 | 307.04 |  | 307.04 | 48.44 | 48.44 |  |  |  |  |  |
| Cayo Agua | 9.155, -82.037 |  | 51.15 |  |  |  |  |  |  |  |  |
| Rio Luis | 8.598, -81.206 | 401.03 | 142.54 |  |  | 142.54 |  |  |  |  | 142.54 |
| Santa Fe | 8.56645, -81.191 | 402.69 | 144.41 | 402.69 | 144.41 |  |  | 109.76 |  |  |  |
| Coclesito | 8.77532, -80.54823 | 473.38 |  |  |  | 214.93 | 214.93 |  | 221.83 |  | 214.93 |
| El Copé | 8.6697, -80.593 |  |  |  | 210.33 |  |  |  |  | 209.87 |  |
| El Valle | 8.633, -80.155 |  |  |  |  |  |  |  |  | 263.49 |  |
| Achiote | 9.18352, -79.98356 | 535.49 |  | 535.49 | 276.65 | 276.65 | 276.65 |  | 283.68 |  | 276.65 |
| Agua Claras | 9.187, -79.69 | 567.63 |  |  |  |  |  |  |  |  |  |
| Gamboa | 9.16933, -79.7529 |  |  |  |  | 302.01 | 302.01 |  |  |  |  |
| Cerro Azul | 9.1611, -79.416 |  | 340.04 |  | 340.04 |  | 340.04 | 304.96 |  | 339.30 | 340.04 |
| Cerro Jefe | 9.2333, -79.35 |  |  |  | 347.34 |  |  | 312.25 |  |  |  |
| Palenque | 9.5734, -79.352 |  | 347.10 |  |  |  |  |  | 353.07 |  |  |
| Upper Rio Chagres | 9.3875, -79.34317 | 605.93 |  |  |  |  | 348.09 |  |  | 347.34 |  |
| San Blas | 9.3558, -78.97929 |  | 388.21 |  |  |  |  |  |  |  |  |
| Lago Bayano | 9.1564, -78.698 |  |  |  | 419.17 |  |  |  |  | 418.26 | 419.17 |
| Cerro Chucanti | 8.78932, -78.45137 | 704.02 | 445.52 |  | 445.52 | 445.52 | 445.52 | 411.09 |  |  |  |
| Tropic Star Lodge | 7.57, -78.19 |  |  |  |  |  | 475.21 |  |  |  |  |
| Aruza Abajo | 8.3613, -77. |  |  |  |  |  |  |  |  | 500.64 |  |
| Puerto Piña | 7.6333, -78.183 |  | 475.95 |  |  |  |  |  |  |  |  |
| El Salto | 8.3135, -77.7884 |  |  |  |  |  | 519.48 |  |  |  |  |
| Rancho Frío | 8.02, -77.732 |  | 525.73 |  |  | 525.73 | 525.73 | 490.25 |  |  |  |
| Cana | 8.02, -77.733 | 783.18 | 525.73 |  | 525.73 |  | 525.73 |  | 531.15 | 524.58 | 525.73 |
| Cemaco | 8.25343, -77.72 |  |  |  |  |  | 527.51 |  | 532.92 | 526.58 |  |

**Table S3:** Regression models of mtDNA pairwise divergence vs other parameters. Models where the regression is significant in bold.

| **Variable** | **Slope** | **Adjusted R^2^** | **p-value** |
| --- | --- | --- | --- |
| Cline width | 6.593 | -0.1093 | 0.7451 |
| Cline width variance | -3739 | 0.1698 | 0.1304 |
| Cline center variance | -492.9 | 0.1495 | 0.1467 |
| **Proportion total SNPs fixed** | **0.02261** | **0.4231** | **0.02479** |
| **Proportion autosomal SNPs fixed** | **0.2266** | **0.4679** | **0.01746** |
| Proportion Z SNPs fixed | 0.03267 | 0.1058 | 0.1887 |
| Enrichment of fixed Z loci | -0.1811 | -0.08119 | 0.5847 |

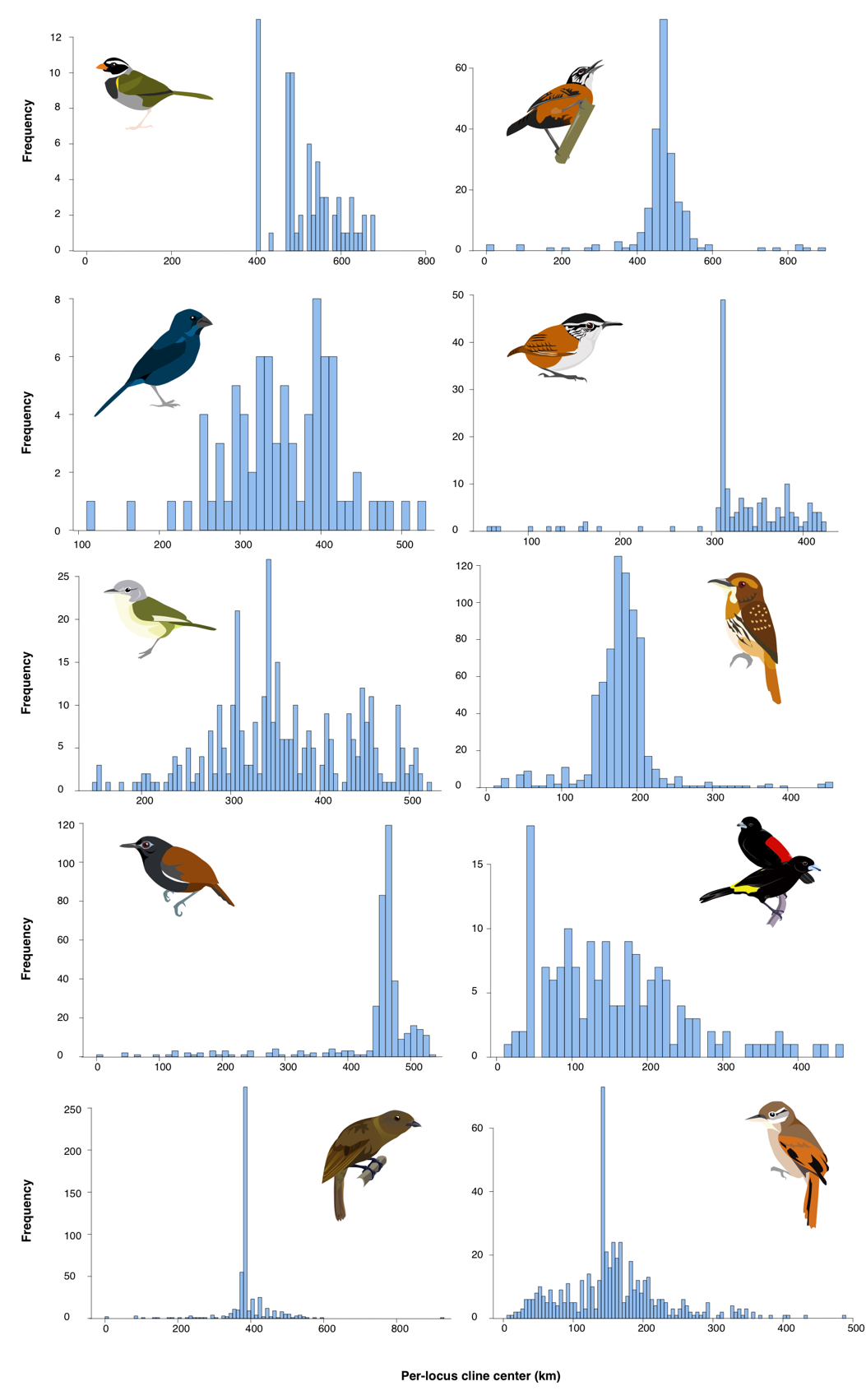

**Figure S1:** Distribution of per-locus geographic cline center estimates fit in HZAR, shown for each study taxon.

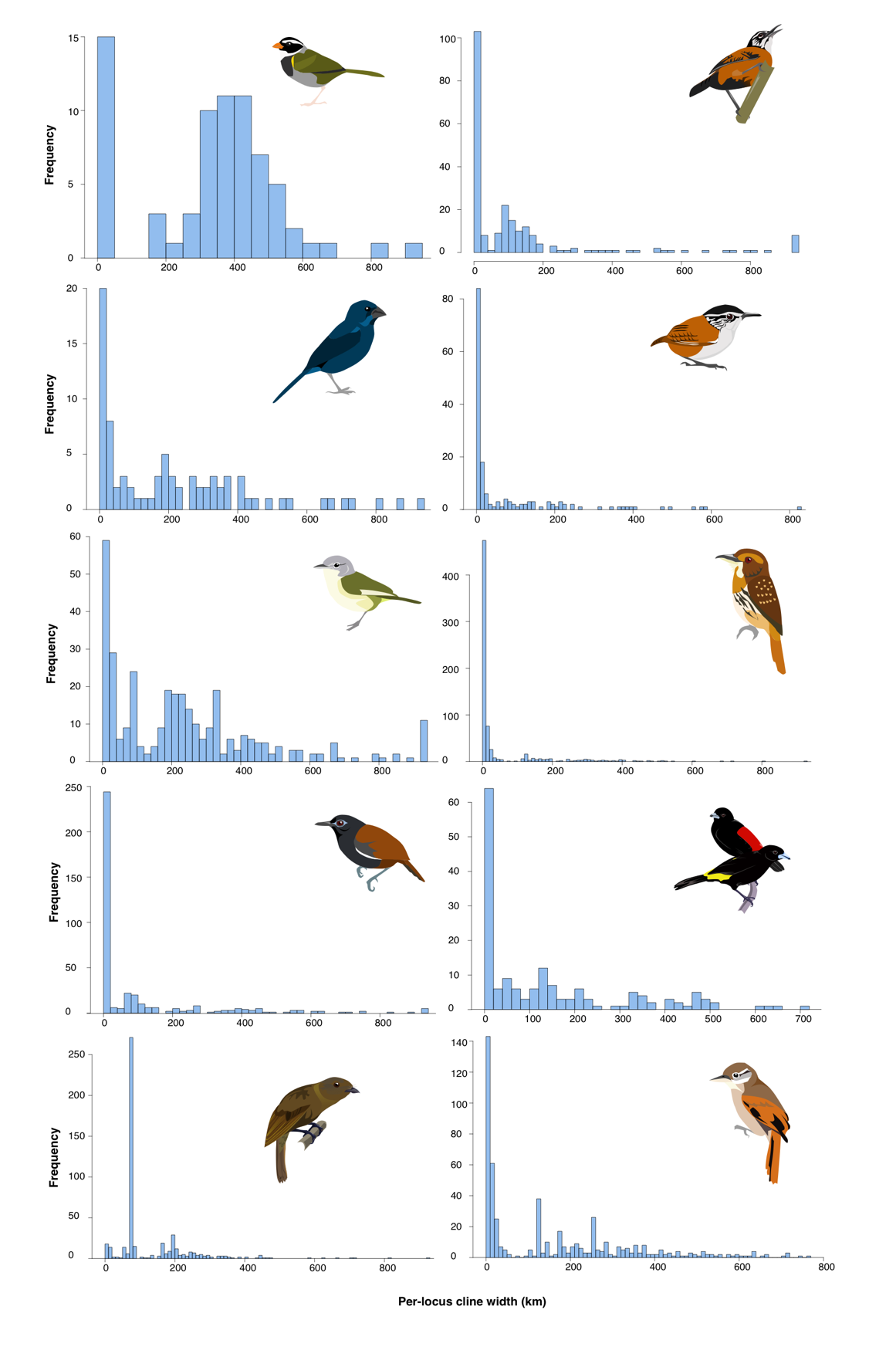

**Figure S2:** Distribution of per-locus geographic cline width estimates fit in HZAR, shown for each study taxon.
